# The circadian gene Dec2 promotes pancreatic cancer dormancy by regulating tumor cell antigen presentation to facilitate immune evasion

**DOI:** 10.1101/2024.11.12.623005

**Authors:** Chris R. Harris, Lan Wang, Crissy Dudgeon, Orjola Prela, Juliana Cazarin de Menezes, Ching-Hua Shih, Christina Davidson, Anthony Casabianca, Subhajyoti De, Wade Narrow, Jennifer Becker, Vinod P. Balachandran, Paul M Grandgenett, Michael A. Hollingsworth, Jean L. Grem, Minsoo Kim, Yeonsun Hong, Scott Gerber, Paula M. Vertino, Chongfeng Gao, Anna Repesh, Zachary Klamer, Yansheng Hao, Brian Altman, Brian Haab, Darren R. Carpizo

**Author notes:** Co-first authors (Chris R. Harris, Lan Wang, Crissy Dudgeon).

## Abstract

The mechanisms that regulate cancer dormancy remain poorly understood. Using a mouse model of resectable pancreatic adenocarcinoma (PDAC), we identified Dec2 as a gene that was upregulated in metastatic dormant tumor cells. Deletion of Dec2 from tumor cells substantially increased mouse survival after resection due to an immune-mediated mechanism as the survival benefit was abrogated in immunodeficient conditions. Dec2 promoted immune evasion by repressing multiple components of the MHC-I dependent antigen presentation pathway in tumor cells. Dec2 is a regulator of circadian rhythms, and we found several components of the antigen presentation pathway oscillated in a circadian manner that was lost upon deletion of Dec2. Moreover, T-cell mediated tumor cell killing varied depending on the time of day. We suggest that lowered MHC-I presentation of antigens during rest phase is a natural effect of the circadian clock, which is exploited by Dec2-overexpressing pancreatic tumors to evade the immune system.

## Introduction

Metastatic recurrence following surgery with curative intent is one of the most significant clinical problems in oncology. Latent recurrences are due to tumor cells that have disseminated to distant organ sites that are occult at the time of surgery and remain dormant for a period (sometimes years) before reactivating to form tumors. While this is true for a multitude of solid organ cancers, it is particularly prevalent in patients with pancreatic adenocarcinoma (PDAC) where virtually all patients (>80%) that undergo curative intent surgery suffer from metastatic relapse indicating that virtually all early-stage patients harbor occult disseminated disease. The mechanisms of metastatic tumor cell dormancy are poorly understood in part because of a lack of animal models that recapitulate the biology of the resected patient. These models are challenging because they require survival surgery as well as extended periods of follow up time. Here we developed the first mouse model of early-stage PDAC that recapitulates the oncologic outcomes of resected patients for the study of PDAC dormancy.

We identified the transcription factor Dec2 (*Bhlhe41*) as upregulated in dormant PDAC cells and validated this in human PDAC. Very high levels of Dec2 were sufficient to induce the quiescent phenotype that is characteristic of dormant tumor cells whereas lower levels of Dec2 could be detected in actively growing tumor cells and was insufficient to cause quiescence. There is emerging evidence that the immune system plays a major role in the metastatic outgrowth of dormant tumor cells (Eyles et al., 2010; Krall et al., 2018; Pommier et al., 2018). Interestingly, the overexpression of Dec2 in both actively growing and dormant cells was sufficient to promote immune evasion through the downregulation of multiple components of the MHC-I-dependent antigen presentation pathway in tumor cells. Loss of Dec2 led to restoration of anti-cancer immunity, inhibition of both primary tumor growth and eradication of disseminated tumor cells. This was accomplished by increasing antigen presentation on tumor cells as well as altering the tumor microenvironment from immunosuppressive to immunoreactive.

Dec2 is a transcriptional repressor that is known to function as part of a negative accessory arm of the molecular circadian clock by opposing the activity of the positive-arm transcription factors CLOCK and BMAL1(Kato et al., 2014). We found that Dec2’s regulation of anti-tumor immunity is through its role as a circadian rhythm factor as proteasome activity, cell surface class I MHC (MHC-I), and T cell-dependent tumor cell killing were also under circadian regulation. This is the first report in which MHC-I has been shown to be under circadian regulation, and while there have been other previous reports that PDACs regulate anti-cancer immunity by modulating MHC-I levels in both actively growing and dormant tumor cells (Pommier et al., 2018; Yamamoto et al., 2020), this is the first report in which PDACs are shown to modulate MHC-I by exploiting a normal function of the circadian clock.

## Results

### A novel model of early-stage pancreatic cancer recapitulates outcomes of resected patients to facilitate the study of dormancy

To gain insight into the mechanisms that contribute to pancreatic cancer cell dormancy and recurrence after surgical resection, we sought to develop a mouse model that recapitulates the patterns of recurrence and survival outcomes of resected patients. Ideally, such a model would exhibit spontaneous dissemination prior to surgical resection in an immunocompetent host since emerging evidence has shown that surgery and the wound healing response induces systemic immune changes that can lead to metastatic outgrowth (Krall et al., 2018). To construct this model, we adapted a syngeneic orthotopic model derived from the Pdx-1-Cre; Kras^G12D/+^; p16^Ink4a -/-^ model to establish the cell line Ink4a.1 that expresses luciferase and mCherry from the same promoter, thus allowing for *in vivo* tumor monitoring and downstream lineage tracing (Collisson et al., 2012). One hundred cells were injected into the distal pancreas of FVB mice and allowed to form a primary tumor for three weeks, at which time a distal pancreatectomy/splenectomy (standard surgery for distal PDAC) was performed, (Fig. 1A, SFig. 1A). Luciferase imaging was used to monitor growth of the primary tumor, to confirm the absence of disease 1-2 days postoperatively and then to monitor for recurrence weekly for 5 months (Fig. 1B).

**Figure 1.**
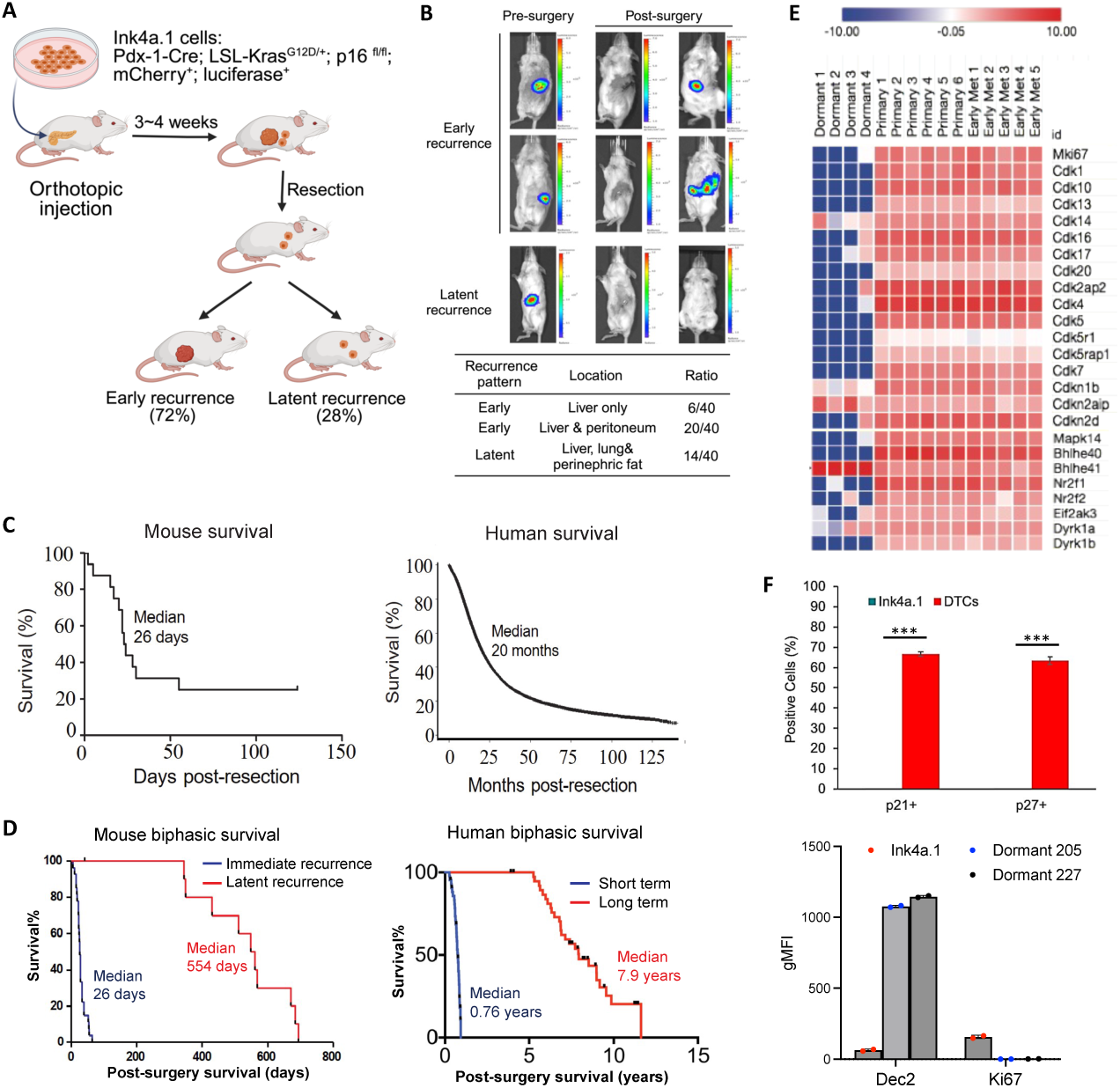
A novel model of early-stage pancreatic cancer recapitulates outcomes of resected patients to facilitate the study of dormancy. (A)Ink4a.1 luc^+^ mcherry^+^ cells were injected into mouse distal pancreas. 3-4 weeks post-implantation, mice underwent distal pancreatectomy with splenectomy and were followed for recurrence. (B)Mice were imaged using the IVIS spectrum following I.P. injection of luciferin for tumor detection prior to resection (left) and 3 days post-resection (middle). The same mice were imaged (right) at 23 days (top), 17 days (middle), and 436 days (bottom) post-surgery. Bar graphs indicate the range of radiance (photons/sec/cm2). Table: The ratio of early to latent recurrent and site of recurrence. (C)Overall survival mouse (left, n=40) and human (right, n= 49,555) following primary tumor resection. Human data was acquired from the National Cancer Database for all resected PDAC patients with T1-3, N0-1 cancers. (D)Mouse and human survival segregated into early and latent recurrence groups. The median survival of mice from the immediate recurrence group was 26 days and the latent recurrence group was 554 days post-surgery (left, n=28 and 12, respectively). Survival of short- and long-term survivors of surgically resected pancreatic ductal adenocarcinoma patients in the MSKCC clinical cohort (right, n=39 and 43, respectively). (E)Heatmap analysis from ultralow RNA-seq analysis of individual dormant DTCs (dormant 1-4) compared with primary tumors (Primary 1-6), and early liver metastases (Early Met 1-5). (F)Flow cytometry analysis of Ink4a.1 cell line or dormant DTCs isolated from livers. Cdk inhibitors p21 and p27 were highly expressed in the dormant DTCs but not in the cell line (n=3/group, top). Dec2 was upregulated and Ki67 was downregulated in dormant DTCs isolated from 2 mice indicated by geometric mean fluorescence intensity (gMFI) level, whereas the primary cell line showed the opposite (bottom). Each black line represents a human or mouse subject (D). Each circle represents a biological replicate with bars indicating mean and error bars indicating standard deviation (SD) (F, bottom). Statistical difference was determined to be p<0.0001 by unpaired t test (F, top).

The two most common distant sites of pancreatic cancer recurrence in humans are the liver and peritoneal cavity, with isolated liver metastases occurring in approximately 25% of cases (Groot et al., 2018; Hishinuma et al., 2006). We found a similar frequency in our model (Fig. 1B), with 15% (n=6/40) of mice exhibiting liver-only recurrence and with 50% (n=20/40) exhibiting both liver and peritoneal recurrence, which occurred early after surgery (mean time to recurrence: 26 days). We found 35% of animals (n=14/40) experienced latent recurrence with a mean time to recurrence of 554 days (Fig. 1B, bottom table). Immunostaining for mCherry in the latent recurrent tumors confirmed they arose from the resected primary tumor (SFig. 1B). For 10/14 mice in the latent recurrent group, we obtained data on sites of recurrence which included the liver, lung, perinephric fat, and uterine lining (SFig. 1C). Kaplan-Meier (KM) overall survival curves for the mice exhibited striking similarity to the survival of early-stage patients after undergoing surgery. There was a steep decline in survival early after resection that leveled out by approximately 65 days with 14 mice (35%) surviving beyond 100 days. Median survival was 26 days (Fig. 1C, left panel), which is equivalent to approximately 27 months in humans (J. G. Fox, 2007). In comparison, we examined the overall survival amongst 41,552 stage I and II PDAC patients who underwent surgical resection (data from the National Cancer Database) and observed similar characteristics with an early decline within the first 20 months that leveled out by approximately 50 months, and a median overall survival of 20 months (Fig. 1C, right panel). Similar outcomes have been published from large single institutional databases (Groot et al., 2018; Winter et al., 2012). This characteristic “L” shape of the KM overall survival curve has not changed in over thirty years in resected PDAC and can be subdivided into two groups: an early recurrent (sharp decline) and latent recurrent (plateau) groups (Winter et al., 2012). In the mouse model we observed the median survival of the early recurrent group to be 26 days whereas that of the latent recurrent group to be 554 days (Fig. 1D, left panel). This appeared similar to a cohort of 82 patients who had undergone surgery for PDAC and stratified into short term (survival 0.25-1 year, median 0.76 years) and long term (survival >3 years, median 7.9 years) survival groups (p<0.0001, Fig. 1D, right panel) (*11*). We conclude that this model recapitulates the outcomes of human early-stage patients undergoing resection for PDAC in both the location and frequency of distant relapse as well as overall survival.

Our model identified a subset of mice exhibiting a long latency period before recurrence. We therefore sought to determine whether we could isolate disseminated tumors cells (DTCs) from the livers of resected, latent recurrent mice to study tumor cell dormancy. Given the rarity of this cell population in the liver, it was important to first validate that we could isolate these cells and verify they originated from the primary tumor. We used fluorescence activated cell sorting (FACS) to sort mCherry^+^ cells (SFig. 1D) from the liver of a mouse with a DFI>200 days (whole body MRI confirmed absence of radiographically detectable disease, SFig.1E). We isolated genomic DNA from the sorted mCherry^+^ cells as well as several controls and PCR amplified with primers specific for luciferase, Kras^WT^, or Kras^G12D^. The luciferase gene was present in the sorted mCherry^+^ cells from this mouse as well as in the unsorted, cultured parental Ink4a.1 cell line, and sorted mCherry^+^ cells from mouse that received an injection of 1×10^6^ cells intrasplenically. Liver DNA from an uninjected mouse was used as a negative control (SFig. 1F). We also detected Kras^G12D^ specific DNA in the sorted mCherry^+^ cells from a mouse with DFI>200 days, as well as the Ink4a.1 cell line and Kras^G12D/+^ tail DNA, but not in the uninjected liver, KRAS^WT^ tail DNA, or no DNA sample (SFig. 1G). These data indicate that the mCherry+ cells isolated from the latent recurrent mice stem from the primary tumor.

To determine if the DTCs from latent recurrent mice exhibited features associated with quiescence we performed RNAseq using an ultralow input protocol. We sequenced three individual sorted cells (Dormant 1-3) as well as a pool of 100 cells (Dormant 4). We examined the expression of markers of proliferation including Ki67 and a large panel of cyclins and cyclin dependent kinases. We found that expression of proliferation genes was decreased in the DTC populations relative to cells from the primary tumor or early recurrent liver metastases (Fig. 1E). Others have shown that the Cyclin-Dependent Kinase inhibitors p21 (Cdkn1a), p27 (*Cdkn1b*), are upregulated in dormant DTCs from other models (Bragado et al., 2013; Sosa et al., 2015). We found that p21 and p27 were similarly upregulated in DTCs relative to primary tumor or early metastases (Fig. 1F, upper panel). Taken together these data suggest that the DTCs obtained from livers of latent recurring mice are in a quiescent state.

Recently several transcription factors have been identified to regulate dormancy including Dec2 (Adam et al., 2009; Sosa et al., 2015). In Fig. 1E, we show that Dec2 (identified as its alias, Bhlhe41) was significantly upregulated in our DTC samples as compared to the primary tumor (Fig. 1E). Several other dormancy markers including Nr2f1,Eif2ak3 (Pommier et al., 2018), Dyrk1a and Dyrk1b were not upregulated (Litovchick et al., 2011). We validated Dec2 expression in dormant DTCs at the protein level by flow cytometry. DTCs isolated from the livers of two latent recurrent mice (205, 227) exhibited significantly increased Dec2 and decreased Ki67 levels compared with the parental Ink4a.1 cells (Fig. 1F, bottom panel). We conclude that DTCs isolated from latent recurrent mice are dormant and can be identified by their increased expression of Dec2.

### Dec2 is a dormancy biomarker in human PDAC

We next sought to determine whether Dec2 might serve as a marker of dormant tumor cells in human PDAC by focusing on the liver as this is the most common site of PDAC metastasis. We examined liver tissue harvested from stage IV patients (n=10) with liver metastases that died of pancreatic cancer as part of a rapid autopsy program. With the assistance of a GI pathologist, we selected areas of liver with obvious metastatic lesions as well as areas that were equivocal for metastatic lesions. The identification of metastatic cancer cells was facilitated by multiplexed glycan immunofluorescence as our previous research showed that the glycan biomarkers CA19-9 and sTRA identifies PDAC cells in the primary tumor with high sensitivity and specificity (Barnett et al., 2017; Gao et al., 2021; Staal et al., 2019). As a control, multiplexed glycan detection of non-metastatic, normal liver tissue showed no sTRA positivity and rare CA19-9 positivity in small bile ductal cells (SFig. 2). In contrast, detection of the liver sections showed regions of interest (ROIs) with positive signals from the metastatic cancer cells. The lesions were marked by either sTRA, CA19-9, or both (SFig. 3A; left, middle, right panels). Each cell was classified as positive or negative for each glycan using a signal intensity cutoff value, and these were used to identify ROIs of glycan-positive cells (SFig. 4). We observed that all 10 of the metastatic liver sections showed positive ROIs (Table S1). These results indicate that the combined sTRA and CA19-9 glycans identify PDAC cells with high sensitivity and specificity both in primary tumors and in the metastatic liver. In addition, we found that the glycan-positive cells appeared both in the large, established cancer lesions (large ROI >1000 cells) as well as in small ROIs (100-1000 cells) and areas that were equivocal by histomorphology (equivocal ROIs, <100 cells, SFig. 5). The proportion of cells that were CA19-9+/sTRA- differed by size of ROI (SFig. 3B, left panel); the proportion of cells CA19-9-/sTRA+ were enriched in the equivocal ROIs with <100 cells as compared to the small or large ROI’s (p<0.0001, mixed effects logistic regression with post hoc comparisons) (SFig. 3B, right panel). Interestingly, previous reports have shown an association between sTRA-exclusive PDAC cells, but not the CA19-9-exclusive cells, with mesenchymal morphology and poor differentiation (Barnett et al., 2017; Gao et al., 2021). These data suggest that the small glycan-positive regions indicate incipient seeding of cancer cells of mesenchymal-type cells, which are not readily observable by imaging or surgical pathology.

Using the glycan-identified ROIs of cells as indicators of metastatic lesions, we analyzed the expression of DEC2 and Ki67 by multiplexed immunofluorescence in the same sections used for the glycan detection. We analyzed a total of 40 large, 129 small and 92 equivocal ROIs. Visual evaluation of selected regions indicated that areas of large ROIs were generally DEC2 low and Ki67 high (Fig. 2A left panel), whereas areas of small and equivocal ROIs were generally DEC2 high and Ki67 low (Fig. 2A middle, right panels). Quantification across cells by size cluster revealed the proportion of Dec2+ cells was significantly higher in the equivocal ROIs compared to both the small and large ROIs (<100 cells vs. >1000 p<0.001; <100 cells vs. 100-1000 p<0.001;>1000 vs. 100-1000 p=0.037) (Fig. 2B). Our evaluation of the relationship between DEC2 and Ki67 showed rare co-expression of DEC2 and Ki67 (Fig. 2C, left panel). Of the total number of cells that were Dec2+, 91% were Ki67-, and of the Ki67+ cells 98% were Dec2-(Fig 2C, right panel). The proportion of cells with high Ki67 was significantly lower in the DEC2-high cells than in the DEC2-low cells (p<0.0001, mixed effects logistic regression with post hoc comparisons) (Fig. 2D), consistent in both the small and large clusters. These findings support that sTRA glycan staining can be used to identify disseminated human PDAC cells in the liver that are equivocal by histomorphology and that Dec2 is a dormancy biomarker in human PDAC.

**Figure 2.**
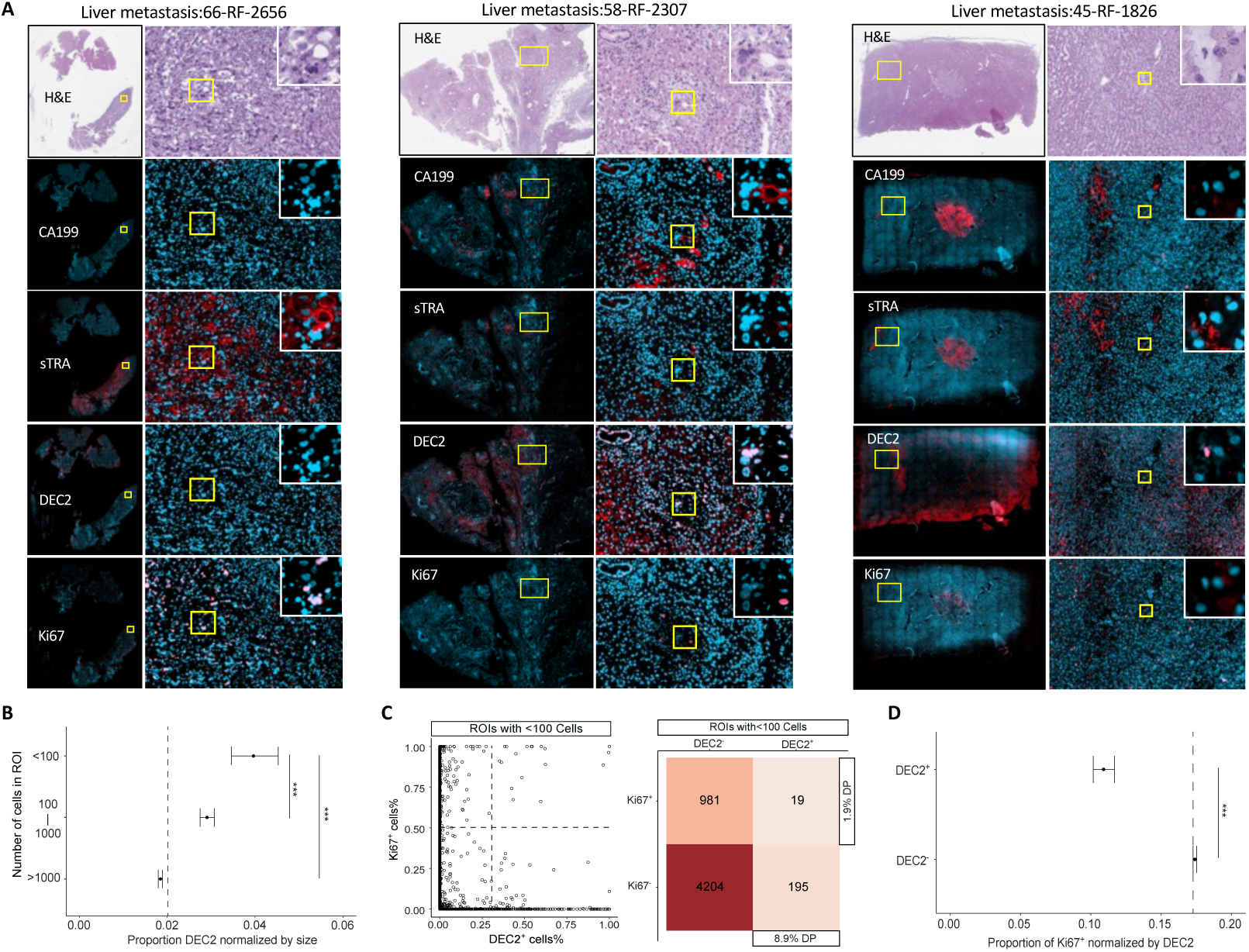
Dec2 is a dormancy biomarker in human PDAC. (A)H&E and multiplexed immunofluorescence images of the PDAC-associated glycans CA19-9 and sTRA, Dec2, and Ki67. Representative images from 3 different cases are shown in which the PDAC cells are positive for sTRA without CA19-9 (left), CA19-9 without sTRA (middle), or both (right). The blue color indicates the nuclear stain DAPI, and the red color indicates each of the markers. Pink indicates an overlap of the red and blue. (B)The plot shows the average and 95% CI of cells that are positive for Dec2 in ROIs composed of <100, 100-1000, or >1000 cells. (C)Scatter plot of the cellular staining of Ki67 with respect to Dec2, where each data point represents a cell among the ROIs composed of <100 cells. The matrix gives the numbers of cells in each quadrant of the scatter plot. (D)The plot gives the average and 95% CI of cells that are positive for Ki67 among either Dec2-positive or Dec2-negative cells in the ROIs. Dash lines indicate the median of positive staining (B-D). Statistical differences were identified by mixed effects logistic regression with post hoc comparisons, with significance against control group indicated by “***” (p<0.0001) (B&D).

### Loss of Dec2 inhibits tumor growth and progression *in vivo* through an immune mediated mechanism

To investigate the mechanistic role of Dec2 in cancer dormancy, we first overexpressed it in the Ink4a.1 cells. Previous reports found that Dec2 promotes cellular quiescence possibly through repression of CDK4 (Adam et al., 2009; Bragado et al., 2013). Consistent with this, we found overexpression of Dec2 in vitro led to a decrease in Ki67 expression (SFig. 6A). To determine if Dec2 regulates *Cdk4*, we constructed a *Cdk4* promoter luciferase reporter in the Ink4a.1 cells. Dec2 overexpression resulted in a decrease in luciferase activity, indicating that Dec2 represses transcription from the *Cdk4* promoter (SFig. 6B). Interestingly we could detect Dec2 in proliferating pancreatic cancer cells *in vitro* suggesting that moderate levels of Dec2 were insufficient to inhibit proliferation (SFig. 6C). We then knocked out Dec2 from the Ink4a.1 cells as well as the 6419c5, which is another murine PDAC cell line derived from the Pdx-1-Cre; Kras^G12D/+^; Trp53^+/-^ model expressing the YFP lineage tracer (KPCY), 6419c5 (Li et al., 2018). We observed a very mild *decrease* in proliferation supporting that moderate levels of Dec2 do not suppress cell proliferation (SFig. 6C,D). We also examined Dec2 mRNA using publicly available data from PDAC tumors compared to healthy pancreas tissues, (Bartha and Gyorffy, 2021) and found higher Dec2 expression in PDAC tumors which is not consistent with an *in vivo* mechanism strictly of inhibiting tumor cell proliferation (SFig. 6E).

To test for a role of Dec2 in actively growing PDAC cells, we implanted Ink4a.1 Dec2 KO cells in the orthotopic resection model. Surprisingly, we found loss of Dec2 slowed primary tumor growth, with the size of Ink4a.1 Dec2 KO tumors being significantly smaller at the time of resection (SFig. 7A, left panel). When we followed the mice after surgery, we found those implanted with Dec2 KO Ink4a.1 cells exhibited a striking improvement in overall survival with a median survival in the Dec2 WT cells of 193 days, while that of mice implanted with the Dec KO cells was not reached (Fig. 3A). These results were not consistent with a tumor suppressive function by Dec2. We hypothesized that the increased survival of the Dec2 KO mice was due to a decrease in the disseminated tumor cell burden. To measure the disseminated tumor cell burden, we performed qPCR for luciferase from genomic DNA harvested from livers of mice that were implanted with Ink4a.1WT or Ink4a.1 Dec2 KO cells and sacrificed after 20 days. We detected markedly diminished luciferase DNA in the livers of mice implanted with Dec2 KO cells as compared to Dec2 WT cells (Fig. 3B) indicating a striking decrease in the disseminated tumor cell burden.

**Figure 3.**
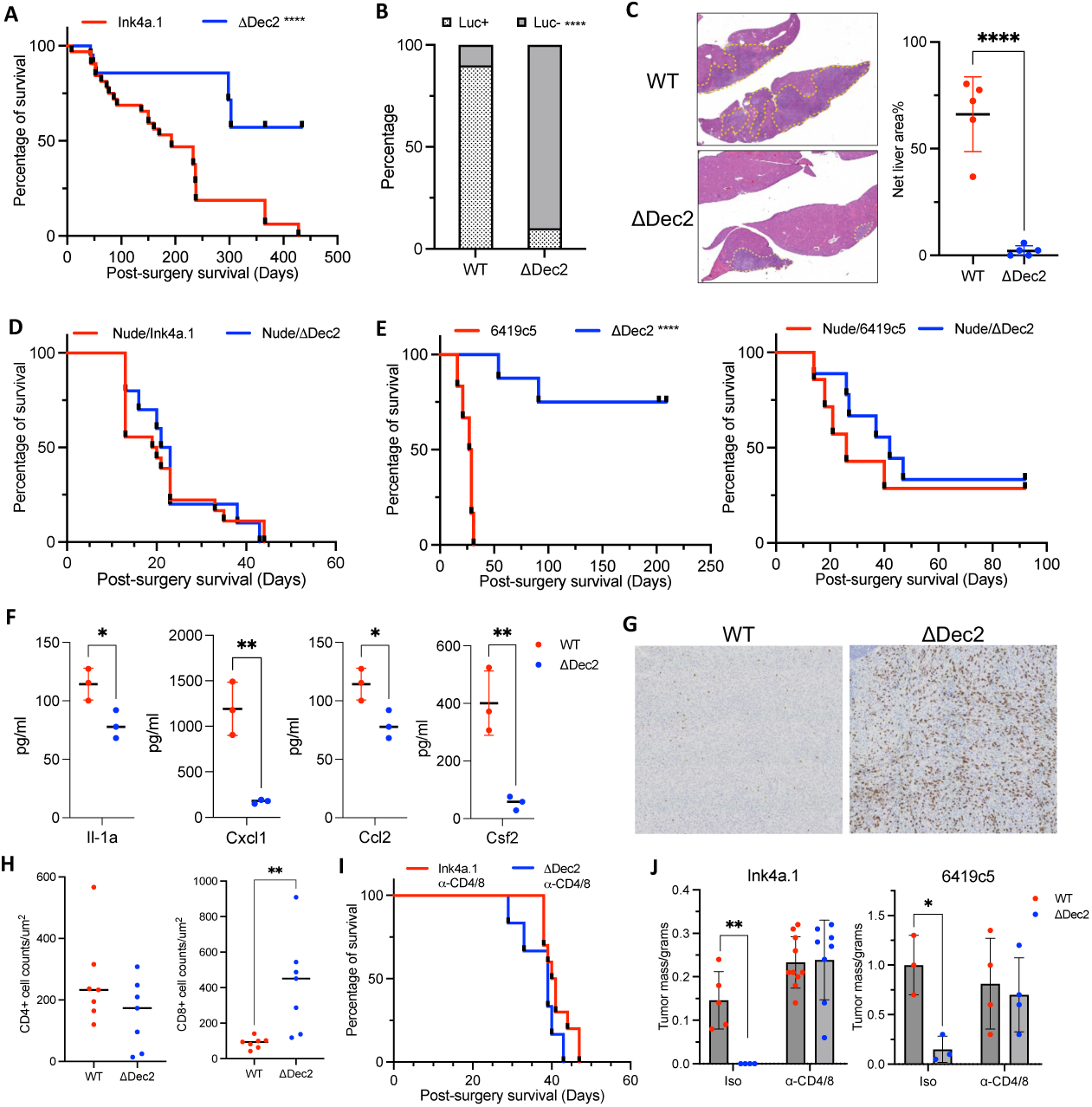
Loss of Dec2 inhibits tumor growth and progression *in vivo* through an immune mediated mechanism. (A)Overall survival of FVB mice orthotopically receiving Ink4a.1 parental (red, n=32) or ΔDec2 (blue, n=21) after primary tumor resection. (B)Liver DTC burdens are determined by tumor-specific luciferase gene quantitative PCR using whole liver gDNA isolated from mice resected for Ink4a.1 WT or ΔDec2 primary tumors (n=10/group). (C)Splenic metastasis assay showed decreased liver metastasis when Ink4a.1 ΔDec2 cells were seeded into the liver compared to the parentals (n=5/group). (D)Overall survival of nude mice receiving Ink4a.1 WT (n=18) or ΔDec2 (n=10) after primary tumor resection. (E)Overall survival of B6 (left) or nude mice (right) receiving 6419c5 WT (red, n=6 each) or ΔDec2 (blue, n=8 and 6). (F)Luminex analysis of primary tumor lysate generating from C57BL/6J mice receiving 6419c5 WT or ΔDec2 (n=3/group). (G)Representative IHC staining of CD8 in 6419c5 WT or ΔDec2 tumor. (H)Quantification of CD4 or CD8 IHC staining on 6419c5 tumors (n=7/group). (I)Overall survival of FVB mice I.P. injected for ⍺-CD4 and ⍺-CD8 *InVivoMab* receiving Ink4a.1 WT (n=10) or ΔDec2 cells (n=6). (J)Tumor mass comparison when mice were treated with isotype or ⍺-CD4 and ⍺-CD8 *InVivoMab*. Each black line or circle represents a single mouse sample (A, C-F, H-J), with horizontal lines/bars indicating mean and error bars indicating SD (C, F, H, J). Statistical differences were identified by log-rank test for Kaplan-Meier curve (A, D&E, I), Fisher’s exact test (B) or unpaired t test (C, F, H, J), with significance against control group indicated by “*” (p<0.05), “**” (p<0.01) or “****” (p<0.001).

It is possible that the decrease in disseminated tumor cell burden is a consequence of the reduction in primary tumor burden observed at resection in the Dec2 KO setting. To further evaluate the impact of Dec2 KO on tumor cell dissemination, we employed a liver metastasis assay where cells were injected into the spleen then immediately splenectomized. After three weeks, the livers were harvested for quantitation of tumor burden. We found that in the mice engrafted with Ink4a.1 Dec2 WT cells over 60% of the liver area was occupied by metastatic tumor versus less than 1% in the Dec2 KO group (Fig. 3C).

The previous experiments were performed in immune-competent FVB mice, and we wondered if the differences in primary tumor growth and survival observed between Dec2 WT and Dec2 KO groups might be through an immune mediated mechanism. To test this, we replaced the immunocompetent FVB with immunodeficient (nude) hosts in the resection model and implanted the Ink4a.1 Dec2 WT and KO cells. Strikingly, we found there was no longer a significant difference in primary tumor size (SFig. 7B right panel), nor was there a significant difference in survival between the two (Fig. 3D). To generalize our findings to other PDAC models, we validated this result in a separate resection model using the 6419c5 cells implanted into immunocompetent B6 mice. Like the Ink4a.1 resection model, we found that loss of Dec2 resulted in smaller primary tumors (SFig. 7C, left panel). Again, this result was abrogated when the cells were implanted in immunodeficient mice (SFig. 7C, right panel). Also similar to the Ink4a.1 resection model, mice implanted with 6419c5 Dec2 KO cells had a strikingly better survival than mice implanted with Dec2 WT cells (Fig. 3E, left panel), a difference that was also abrogated when the cells were implanted in nude mice (Fig. 3E, right panel). Taken together, we conclude that Dec2 promotes tumor progression through the suppression of anti-tumor immunity.

We next characterized the tumor immune microenvironments of orthotopic tumors from WT and Dec2 KO cells. 6419c5 is often used for this purpose because it is known to generate PDACs expressing elevated levels of immunosuppressive cytokines such as Cxcl1 that lead to poor T-cell trafficking, a condition that is often called immunologically cold (Li et al., 2018). We examined the cytokine profiles of 6419c5 Dec2 WT and KO tumor tissue using a Luminex array and detected significant decreases in a number of cytokines known to promote an immunosuppressive microenvironment in PDAC such as Cxcl1, IL1a, IL-6, Ccl2 and Csf2 (GM-CSF) in the Dec2 KO tumors relative to WT (Fig. 3F) (Bayne et al., 2012; Bianchi et al., 2023; Li et al., 2018; Sanford et al., 2013; Singh et al., 2024) . Except for IL-1a, these cytokine changes *in vivo* were not due to a change in gene expression in the tumor cells as we observed an increase in Cxcl1, Ccl2 and Csf2 upon Dec2 loss *in vitro* (SFig. 7C). We examined both CD4 and CD8 T-cell populations by immunohistochemistry and flow cytometry in the 6419c5 Dec2 WT and KO tumors and found there was no significant difference in CD4 cells; however, CD8 T-cells were significantly increased in the Dec2 KO tumors (Fig. 3G,H). Interestingly we found in the Ink4a.1 model that loss of Dec2 led to a significant increase in CD4 T-cells while CD8 T-cells were not significantly increased (SFig. 7D). We next treated immunocompetent FVB mice in the Ink4a.1 resection model with anti-CD4 and CD8 antibodies to immunodeplete T-cells and found that depletion of CD4 and CD8 T cells abrogated the survival advantage conferred by Dec2 KO (Fig. 3I). CD4 and CD8 T-cell depletion also abrogated the primary tumor growth inhibition observed with loss of Dec2 in both Ink4a.1 and 6419c5 tumors (Fig. 3J). Taken together these results indicate that Dec2 deletion results in restoration of anti-tumor immunity and repolarization of the immune microenvironment from suppressive to active through a T-cell dependent mechanism. We also conclude that the effect of Dec2 on tumor cells depends on the level of expression where high levels inhibit cell proliferation and anti-tumor immunity whereas moderate levels allow cell proliferation but inhibit anti-tumor immunity and low or absent levels allow cell proliferation and immune recognition.

### Dec2 regulates multiple components of the antigen presentation pathway

To investigate the role of Dec2 in antitumor immunity, we performed a differential expression analysis of bulk RNAseq data from 6419C5 and Ink4a.1 WT and Dec2 KO cells (SFig. 8A). Given the effect of T-cells on tumor growth and survival in mice injected with WT vs Dec2 KO cell lines, we were particularly struck by the fact that Dec2 loss in both cell lines affected genes involved in MHC-I dependent antigen display. This is the pathway that produces the antigens recognized by CD8 T-cells.

We used flow cytometry of intact cells to determine whether Dec2 affected the amount of MHC-I on the surfaces of several PDAC cells lines including Ink4a.1 , 6419c5 and A2441 cells (derived from the PDAC model *Ptf1a-Cre; Arid1a^flox/flox^* (Wang et al., 2019)). Ink4a.1 cells expressed higher levels of MHC-I at baseline, whereas 6419c5 cells expressed almost no MHC-I and A2441 cells were intermediate. As shown in Fig. 4A, there was a significant increase in surface MHC-I in the cell lines with moderate to high MHC-I (Fig.4a Ink4a.1 left panel), A2441 (Fig. 4A middle panel) as well as the immunologically “cold” 6419c5 cell line (Fig. 4A right panel). We also knocked down Dec2 in a human pancreatic cancer cell line, BxPC3, and again observed an increase in surface MHC-I (SFig. 8B,C).

**Figure 4.**
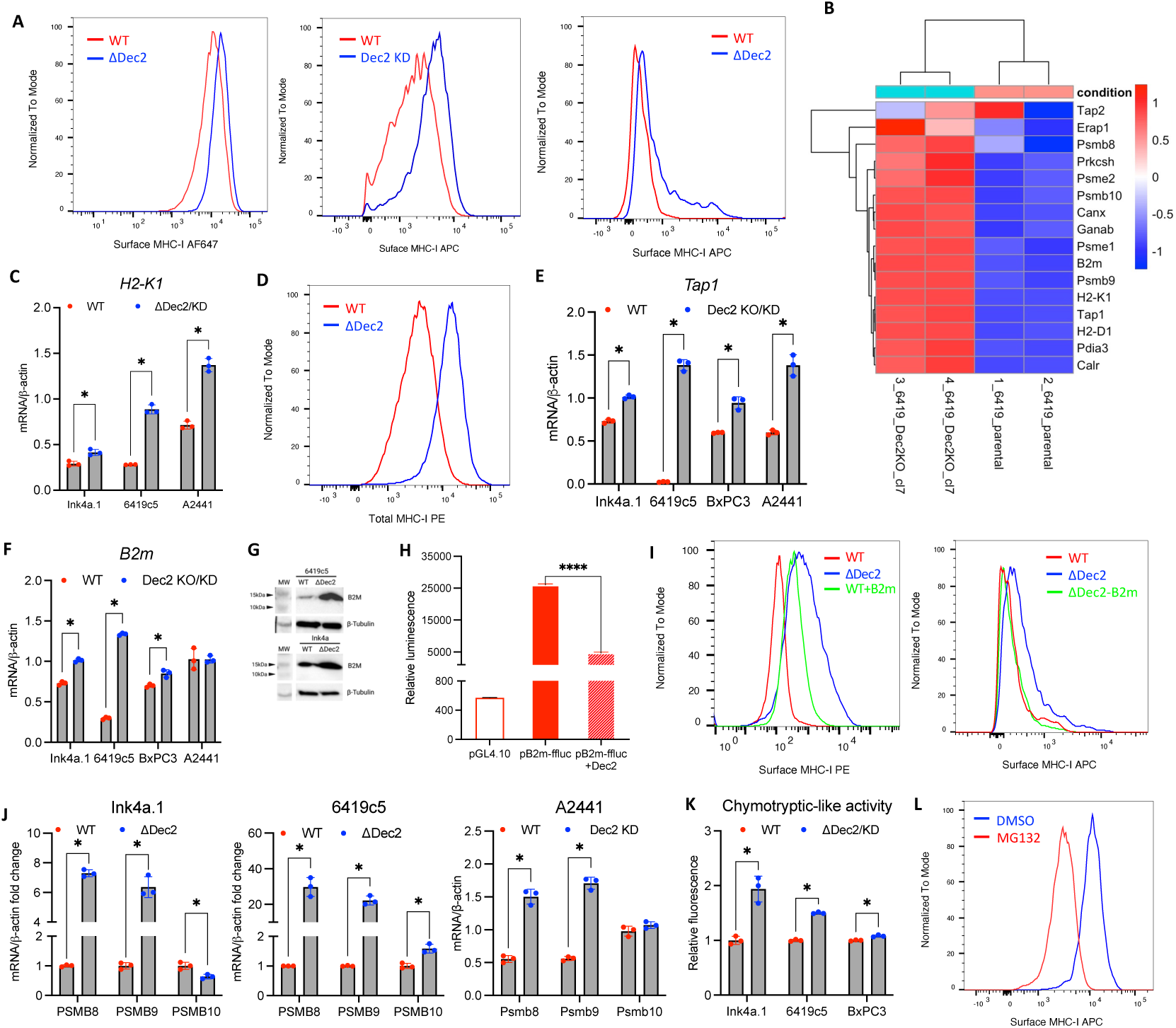
Dec2 regulates multiple components of the antigen presentation pathway. (A)Flow cytometry analysis showed that ΔDec2/KD increased cell surface MHC-I levels in Ink4a.1 (left), A2441 (middle) and 6419c5 (right). (B)Heatmap analysis from bulk RNAseq showed an upregulation of genes involved in MHC-I antigen presentation pathway in 6419c5 ΔDec2 compared to the parental (n=2/group). (C)qPCR analysis showed an increase of *H-2k1* expression in ΔDec2/KD cell lines. (D)Flow cytometry analysis showed an increase of total MHC-I in 6419c5 ΔDec2 compared to the parental. (E)qPCR analysis showed an increase of *Tap1* expression in ΔDec2/KD cells. (F)qPCR analysis showed an increase of *B2m* expression in ΔDec2/KD cells. (G)Western blot analysis showed an increase of B2m protein in Ink4a.1 and 6419c5 ΔDec2 cells. (I)Dec2 represses B2m promoter. When Dec2 is overexpressed, *B2m* promoter driven-firefly luciferase (pB2m-ffluc) expression is significantly downregulated (n=3/group). (I) Flow cytometry analysis showed that overexpression of *B2m* in 6519c5 parental cells increases its surface MHC-I (left), whereas knocking down *B2m* from 6419c5 ΔDec2 decreases its surface MHC-I (right). (J)qPCR analysis showed an increased expression of immunoproteasome subunit genes in ΔDec2/KD cells. (K)ΔDec2/KD led to increased chymotryptic proteosome activity in mouse and human PDAC cell lines. (L)Flow cytometry analysis showed blockage of MHC-I surface localization upon MG132 treatment. Each circle represents a replicate, with horizontal lines/bars indicating mean and error bars indicating SD (C, E, F, H, J, K). Statistical differences were identified by multiple unpair t test (C, E, F, J, K) or unpaired t test (H), with significance against control group indicated by “*” (p<0.05) or “****” (p<0.001).

Production of peptides bound to cell surface MHC-I complexes is a multi-step process, and the RNAseq data pointed to the upregulation of multiple genes within this pathway upon Dec2 loss (Fig. 4B), including the genes encoding polypeptides of the MHC-I complex itself (*H2-K1, H2-D1, B2m*), proteasome and immunoproteasome genes (*Psme1, Psmb 8, 9,10*), genes involved in peptide transport (*Tap1*) and in proper folding of MHC-I (*Pdia3*). Many of these transcript differences were validated by quantitative PCR in multiple pancreatic cell lines. For instance, we found *H2-k1* levels to be significantly increased in Ink4a.1, 6419c5 and A2441 cells. Using Cleavage Under Targets & Release Using Nuclease (Cut&Run) comparisons of DNA from Dec2 WT and KO cells, we observed that Dec2 binds to the *H2-k1* promoter in both the Ink4a.1 and 6419c5 cells. This fits with the known biology of Dec2 as a basic-helix-loop helix DNA binding protein whose orange domain represses promoters to which it binds. Dec2 KO in the 6419c5 cells resulted in an increase in H3K27 acetylation, a histone modification associated with open chromatin and a corresponding increase in RNAseq signal relative to WT cells, consistent with gene activation (SFig. 8E, top panel). *H2-d1* mRNA levels were increased in the 6419c5 and A2441 cells but not in in the Ink4a.1 cells (SFig. 8F) and Cut&Run data revealed similar findings to *H2-k1* with evidence of binding of Dec2 at the promoter and in 6419c5 Dec2 KO cells an increase in H3K27 acetylation and a corresponding increase in RNAseq signal relative to WT (SFig. 8E, middle panel). We also examined total cellular MHC-I protein levels in both Ink4a.1 and 6419c5 cells by either flow cytometry (6419c5 cells) or western blot (Ink4a.1 cells) as well as immunofluorescence (both) and detected an increase in the Dec2 KO cells compared to WT controls (Fig. 4D, SFig. 8G,F).

Transporter associated with antigen processing (*Tap1*), which functions to import small peptide antigens processed by the proteasome from the cytosol to the lumen of the endoplasmic reticulum, is significantly upregulated upon Dec2 KO/KD in the Ink4a.1, 6419c5, BxPC3 and A2441 cell lines (Fig. 4E). Cut&Run data from Dec2 wildtype and knockout cells indicated that Dec2 binds to the *Tap1* promoter, and in the setting of Dec2 loss it associates with increased H3K27Ac, decreased H3K27me3 and a corresponding increase in RNAseq signal relative to WT cells, consistent with gene activation (SFig. 9A).

Beta 2-microglobulin (*B2m*) is an integral component of the MHC-I complex. Like *Tap1*, *B2m* mRNA levels were increased relative to controls when Dec2 was knocked out or knocked down in the mouse lines Ink4a.1, 6419c5, BxPC3 well as in A2441 cells (Fig. 4F). B2m protein levels were also upregulated in the Ink4a.1 and 6419c5 Dec2 KO cells (Fig. 4G). Like *Tap1*, Dec2 bound directly to the *B2m* promoter by Cut-and-Run analysis, and upon Dec2 KO it again associated with an increase in H3K27ac and a decrease H3K27me3 (SFig. 8E, bottom panel). To determine if Dec2 had a direct impact on the *B2m* promoter, we created a *B2m* promoter-luciferase reporter construct. Co-transfection of the reporter and a Dec2 expressing plasmid repressed luciferase activity (Fig. 4H). Enforced overexpression of B2m in 6419c5 cells resulted in a robust increase in cell surface MHC-I, like that observed in Dec2 KO cells (Fig. 4I, left panel). Conversely, the increase in MHC-I surface expression observed in Dec2 KO cells was abrogated in the presence of *B2m* siRNA (Fig. 4I, right panel). Taken together, these data suggest that Dec2 represses *B2m* transcription to control cell surface MHC-I.

The ubiquitin-proteasome system is the source of antigenic peptides bound to and presented by MHC-I. Moreover, proteasome activity and specifically immunoproteasome activity is linked to numerous aspects of anti-tumor immunity including activation of inflammation and T-cell differentiation, all of which we observed in Dec2 KO tumors (Ferrington and Gregerson, 2012; Kloetzel and Ossendorp, 2004). As proteasomal degradation was a pathway highly upregulated in 6419c5 Dec2 KO cells (SFig. 8A), we investigated this as potential mechanistic contributor. We found in the Ink4a.1 Dec2 KO cells a marked upregulation of the catalytic subunits of the immunoproteasome Psmb 8 and 9 but not Psmb 10, while in the 6419c5 and A2441 cells we observed an increase in all three (Fig. 4J). Interestingly the Psmb9 gene shares the same promoter region as Tap1 and by Cut&Run analysis, the Psmb9 promoter was also decorated with Dec2 in the WT lines, and demonstrated more activating H3K27ac marks and decreased H3K27me3 marks in the 6419c5 Dec2 KO (SFig. 8E). Catalytic activity of the proteasome and in particular immunoproteasome activity is characterized by an increase in chymotrypsin-like cleavage after hydrophobic amino acids. We examined the chymotrypsin-like activity in several murine and human PDAC cells in which Dec2 was either knocked out or knocked down and found a significant increase (Fig. 4K). To determine if the increase in chymotryptic-like proteosome activity in the Dec2 KO cells contributed to the increase in cell surface MHC-I expression, we treated the Ink4a.1 Dec2 KO cells with MG132, which inhibits the chymotrypsin-like activity of the proteasome, and observed a significant decrease in cell surface MHC-I (Fig. 4L).

Nascent, not-yet-folded cytosolic proteins are rapidly proteolyzed and serve as the source of most of the peptides that become linked to MHC-I complex. We decided to follow one of these nascent cytosolic proteins, firefly luciferase which is expressed in the Ink4a.1 cell line and is a known immunogen of the mouse immune system (Ferrari et al., 2024). Indeed, Ink4a.1 cells that express luciferase form smaller tumors in immunocompetent mice than cells that express no luciferase (SFig. 8J). We used a cycloheximide pulse followed by a short chase to watch the stability of this protein. Nascent luciferase was less stable in the Ink4a.1 Dec2 KO cell line, which has increased proteasomal activity, than in the Ink4a.1 WT cell line, which has decreased proteasomal activity (SFig. 8K). As a foreign protein, presumably the increased turnover of luciferase in the Ink4a.1 Dec2 KO cells results in more display of foreign luciferase antigens by Dec2 KO tumors. Taken together, these data support a model in which Dec2 regulates multiple components of the antigen presentation pathway ranging from the generation of antigenic peptides by the proteasome to the transport of these peptides (*Tap1*) into the endoplasmic reticulum, to the MHC-I complex genes including H2-k1, H2-d1, B2m.

### Dec2 regulates tumor cell antigen presentation through its role as a circadian rhythm gene

Dec2 has been found to regulate the circadian clock in various peripheral tissues such as the liver, where it regulates the oscillation of several sterol metabolizing cytochrome p450 genes (Honma et al., 2002; Noshiro et al., 2004). We wondered if Dec2 is also important for circadian rhythms within pancreatic cancer cells. We performed a time-series qPCR analysis of Ink4a.1 and 6419c5 cells to investigate the overall impact of Dec2 expression on the components of the molecular clock and used the ECHO algorithm to computationally determine circadian oscillations of gene expression (De Los Santos et al., 2020). Ink4a.1 and 6419c5 Dec2 WT and KO cells were treated with dexamethasone to synchronize their circadian clocks (Balsalobre et al., 2000), which corresponds to ‘circadian time’ zero (CT=0). After 24 hours, at 4-hour intervals over the next 48 hours, gene expression levels were normalized to non-circadian housekeeping genes (*Ppia* for Ink4a.1 and *Tbp* for 6419c5 cells) (SFig. 9A,B). The qPCR analysis revealed that both cell lines show robust circadian oscillation of *Bmal1*, *Per2*, *Cry1*, and *Dbp*, key molecular clock components (Fig. 5A, SFig. 9A,B,C). Notable changes in oscillation amplitude and phase of these genes were observed in Dec2 KO cells when compared to WT, however ECHO analysis demonstrated that gene expression oscillation of these components remains circadian in Dec2 KO cells (Fig. 5A, SFig. 9A,B). These data suggest that the absence of Dec2 does not result in a complete loss of rhythmicity of the clock machinery but drives a distinct change in the amplitude of circadian clock components, and thus a potential change in how downstream genes are controlled.

**Figure 5.**
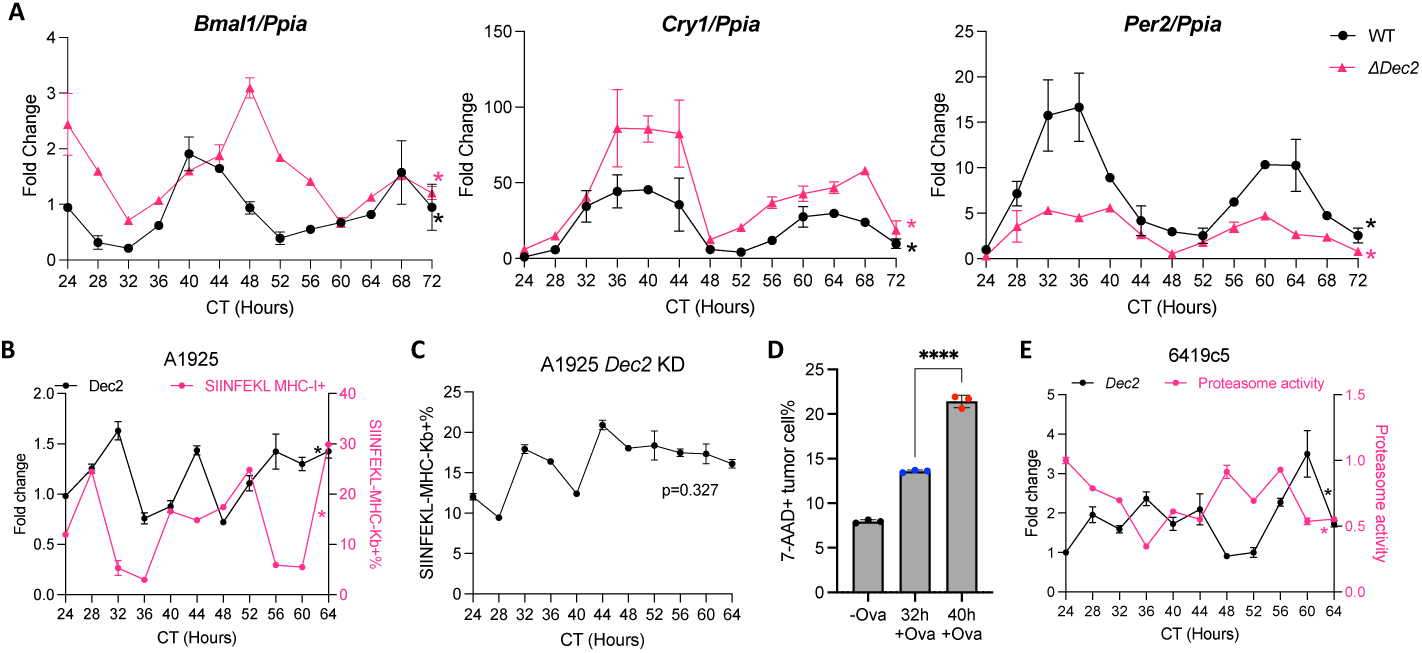
Dec2 regulates tumor cell antigen presentation through its role as a circadian rhythm gene. (A)Circadian regulatory genes in Ink4a.1 WT and ΔDec2 expression by circadian time (CT) from 24 to 72 hrs (n=3/group) when synchronized with dexamethasone. Data was normalized to *Ppia expression*. (B)Flow cytometry analysis of surface MHC-I bound to SIINFEKL peptide (SIINFEKL MHC-I^+^) and qPCR analysis of Dec2 mRNA at matching timepoints (CT 24-64 hrs) of A1925 when synchronized with forskolin. mRNA is normalized to *Rplpo* expression. (C)Flow cytometry analysis of surface SIINFEKL MHC-I^+^ of A1926 *Dec2* KD from CT 24-64 hrs. (D)Quantification of 7-AAD^+^ A1925 cells when co-cultured with CD8^+^ T cells isolated from OT-1 mouse spleen (n=3/group). A1925 were synchronized for 32 h or 40 h and treated with ovalbumin (ova) before co-culture. Cells not treated with ova were used as a control. 7AAD positivity was analyzed by flowcytometry. (E)The chymotryptic proteasomal activity and qPCR analysis of Dec2 mRNA at matching timepoints (CT 24-64 hrs) of 6419c5 when synchronized with dexamethasone. Each circle represents the mean of technical triplicates (A-C, E) or a replicate with bars indicating mean (D). Error bars indicates SD (A-E). Rhythmicity was determined by Extended Circadian Harmonic Oscillator (ECHO) application and indicated by “*” (p<0.01) (A-C, E). Statistical difference was identified by unpaired t test and indicated by “****” (p<0.0001) (E).

Infiltration of T-cells into tumor cells was recently shown to be circadian; this helps to explain why patient response to immunotherapy is strongly affected by the time of day at which the treatment is initiated (Wang et al., 2024). Thus we wondered if MHC class I-dependent presentation of antigens is also circadian. We therefore performed a time-series analysis using A1925 cells, which express MHC-Kb that is used for analysis of binding to the model ovalbumin K^b^-binding peptide SIINFEKL. A1925 cells were synchronized using forskolin instead of dexamethasone as glucocorticoids like dexamethasone have been shown to repress MHC-I expression (Deng et al., 2021). Upon forskolin synchronization, A1925 cells exhibited an intact clock machinery as evidence by *Per2* expression by ECHO analysis (SFig.9D,E). Synchronized A1925 cells were collected at various time points, and to quantitate empty MHC-I complexes on their surfaces, the intact cells were treated with the SIINFEKL peptide, followed by flow cytometry with an antibody that specifically detects SIINFEKL bound to MHC-Kb. Strikingly, ECHO analysis demonstrated that surface MHC-I levels were circadian with the levels anti-phase with Dec2 expression (Fig. 5B). When we evaluated this in A1925 Dec2 KD cells, we found the circadian rhythmicity of MHC-I was lost (Fig. 5C, SFig. 9F).

To determine if the oscillation in surface MHC-I has functional significance, we evaluated antigen specific T-cell killing of the SIINFEKL peptide loaded A1925 cells at two time points (32 and 40 hours) by co-culturing the tumor cells with OT-1 T-cells that recognize SIINFEKL peptide (Hogquist et al., 1994). These time points corresponded to relatively high Dec2 and low MHC-I (CT32 hrs) versus low Dec2 and high MHC-I expression (CT40 hrs), respectively. We observed a significant increase in tumor cell killing from 32 to 40 hours (p<0.001) (Fig. 5D). These results indicate that the circadian clock machinery is intact in pancreatic cancer cells and that antigen presentation and antigen-specific T-cell mediated tumor cell killing is under circadian regulation by Dec2.

Finally, we evaluated whether there is circadian control over other components of antigen presentation. We performed a time-series analysis of proteasome activity in 6419c5 cells synchronized with dexamethasone. ECHO analysis revealed that proteasome activity is also circadian and was highest when Dec2 expression was lowest and vice-versa (Fig. 5E). We also evaluated the expression of genes that generate MHC-I. In 6419c5 WT cell lines, ECHO analysis revealed circadian rhythmicity in the oscillation of and *B2m* and *H2-d1* mRNA levels that was lost upon deletion of Dec2 (SFig. 9G). While the *H2-k1* oscillation in the Dec2 KO cells was still significant by ECHO analysis, a difference in amplitude was evident. Similar results were obtained for *H2-d1* in Ink4a cells, which also showed a loss of rhythmicity in response to Dec2 KO (SFig. 9A).

## Discussion

We have elucidated a novel mechanism by which MHC-I is prevented from displaying tumor antigens in pancreatic cancer. Dormant tumor cells and pancreatic tumor tissue (relative to normal pancreas) expressed elevated levels of Dec2, a DNA binding protein that represses the promoters of several genes involved in assembly of antigen/MHC-I complexes. Dec2 reduced the expression of proteasome proteins that produce the peptides displayed by MHC-I, as well as Tap1 which transports these peptides across the ER membrane, and B2m that is an essential part of the peptide/MHC-I complex.

Pancreatic cancer is characterized by a highly immunosuppressive tumor microenvironment that is thought to be the basis for its resistance to immune checkpoint inhibition(Li et al., 2018; Ullman et al., 2022). This immunosuppressive TME has several mechanisms including an abundance of both granulocytic and monocytic myeloid derived suppressor cells as well as carcinoma-associated fibroblasts (CAFs) which all contribute to inadequate CD8 T-cell infiltration and activation (Bayne et al., 2012; Bianchi et al., 2023; Jiang et al., 2016; Li et al., 2018). Efforts to therapeutically target these populations in the clinic have thus far been unsuccessful (Noel et al., 2020; Padron et al., 2022; Ullman et al., 2022; Wang-Gillam et al., 2022). One reason for this may include insufficient MHC-I levels on the surface of tumor cells.

Low expression of surface MHC-I in PDAC as a mechanism of immune evasion is supported by recent studies demonstrating regulation of MHC-I by other molecular mechanisms such as the ER stress response (Pommier et al., 2018) and autophagy (Yamamoto et al., 2020). The mechanism by which Dec2 loss promotes antitumor immunity is distinct in that it not only increases MHC-I presentation but also repolarizes the immune microenvironment from immunosuppressive to immunoreactive. While further mechanistic studies are needed to dissect exactly how this happens and how it may vary across model systems, one possibility is an increase in proteasome or immunoproteasome activity. PSMB 8,9 and 10 are three major catalytic subunits of the immunoproteasome and increase neoantigen quality by increasing chymotrypsin-like proteasome activity which has been shown to increase cytotoxic T-cell killing and response to immune checkpoint inhibitors in other cancers such as melanoma and non-small cell lung cancer (Javitt et al., 2023; Kalaora et al., 2020). The combination of a mechanism that both increases tumor cell antigen presentation as well as repolarizes the TME makes Dec2 an attractive therapeutic target.

Notably, Dec2 is known to be a regulator of the circadian rhythm, where in humans its expression in the brain increases at night (the rest phase) so that it can repress transcription of Orexin, which encodes a neuropeptide that is responsible for wakefulness and appetite. We now show that Dec2 also causes circadian production of peptide/MHC-I complexes in pancreatic tumor cells. There is now a significant body of evidence in multiple solid organ metastatic cancers supporting greater efficacy of immune checkpoint inhibitor therapy and overall survival when administered in the morning versus the evening hours (Catozzi et al., 2024; Cortellini et al., 2022; Dizman et al., 2023; Karaboue et al., 2022; Qian et al., 2021). Our results provide a tumor cell intrinsic mechanism to support these clinical observations. Since the expression of Dec2 varies across tissue types, our results of the circadian nature of antigen presentation and T-cell mediated tumor cell killing may not apply generally to all tumor cell types.

Our discovery of Dec2’s effect on MHC-I-dependent peptide display in actively growing pancreatic tumors came about through our interest in understanding how dormant pancreatic cancer cells, which live for years within a patient with no evidence of disease, can evade the immune system for such a long period of time. We developed the first mouse model of early stage resected PDAC that mimics human outcomes and discovered that the dormant cells within this model express Dec2 at even higher levels than seen in actively growing tumor cells. It is not clear if dormant tumor cells usurp circadian control of the antigen presentation pathway by chronically expressing Dec2 at high levels or if Dec2 maintains its circadian rhythmicity in metastatic tumor cells. The latter would suggest that metastatic tumor cells might fluctuate in and out of dormancy over time. The elevated amount of Dec2 expressed by dormant cells enables them to evade the immune system by MHC-I repression but may also allow Dec2 to bind to other target promoters, including genes involved in cell proliferation. Indeed, we found that upon overexpression, Dec2 can shut down expression of the critical proliferation gene Cdk4, whose levels are repressed in dormant PDAC cells. Overexpression of Dec2 also caused actively growing PDAC cells to go into the quiescent state characteristic of dormant tumor cells. Thus, the cellular outcome in tumor cells is dictated by the level of Dec2 expression where absent or low levels enable an immune response by the host, moderate levels allow immune evasion without blocking tumor cell proliferation and high amounts block both proliferation and an immune response.

## Acknowledgments

This work was supported by funding from the University of Rochester Medical Center Department of Surgery to DRC and CRH. It was also supported by a pilot grant from the Wilmot Cancer Institute (DRC, PMV, BA). Funding support from the NCI to B.A. (1R01CA282225) and P.M.V. (5R01CA250531). We thank Matthew Raymonda and Josh Munger (University of Rochester Medical Center) for important contributions. We also would like acknowledge assistance from the Wilmot Cancer Institute shared resources in both Genomics and Cytometry. Services, results and/or products in support of the research were generated by Rutgers Cancer Institute of New Jersey Flow Cytometry & Cell Sorting Shared Resource P30CA072720-5921. The following grants provide funding support for the University of Nebraska Medical Center Tissue Bank Rapid Autopsy Program for Pancreas: Pancreatic Cancer Detection Consortium (U01CA210240), NCI Cancer Center Support Grant (P30CA36727), NCI Research Specialist (R50CA211462).

## Author Contributions

1) C.R.H. – investigation, formal analysis, validation, data curation, writing – review, editing
2) L.W. - investigation, formal analysis, validation, data curation, writing
3) C.D. – investigation, formal analysis, validation, data curation, writing – Original Draft, Review and editing
4) O.P. - investigation, formal analysis
5) J.C.M - investigation, formal analysis, writing
6) C.S. - investigation, formal analysis
7) C.D. - investigation, formal analysis
8) A.C. - investigation, formal analysis
9) S.D. - investigation, formal analysis
10) W.N. – investigation, resources
11) J.B. - investigation, formal analysis
12) V.B. - investigation, formal analysis
13) P.G. – resources
14) M.H. – resources
15) J.L.G – resources
16) M.K. – resources
17) Y.H. – investigation
18) S.G. – formal analysis, writing – review, editing
19) P.M.V. - investigation, formal analysis, writing – review, editing
20) C.G. - investigation, formal analysis
21) A.R. - investigation, formal analysis
22) Z.K. - investigation, formal analysis
23) Y.H. – investigation
24) B.A. - investigation, formal analysis, validation, data curation, writing –review, editing
25) B.H. - investigation, formal analysis, validation, data curation, writing –review, editing
26) D.R.C. - investigation, formal analysis, validation, data curation, writing – Original draft, review, editing

## STAR Methods

Key resources table.

Key resources table
| REAGENT or RESOURCE | SOURCE | IDENTIFIER |
| --- | --- | --- |
| Antibodies |  |  |
| TruStain FcX™ (anti-mouse CD16/32) Antibody | Biolegend | 101319 |
| Anti-SHARP1 primary antibody | Abcam | ab175544 |
| Anti-p21 Alexa Fluor® 647 | Abcam | ab237265 |
| anti-mouse p27 Kip1 PE | Cell Signaling Technology | 12184S |
| Anti-mouse/human Ki-67 APC/Cyanine7 | Biolegend | 151231 |
| Anti- human SHARP1 | Abcam | ab244508 |
| Anti-mouse CD44 Alexa Fluor® 700 | Biolegend | 156009 |
| Anti-mouse CD133 PE/Cyanine7 | Biolegend | 141209 |
| APC/Cy7® Conjugation Kit | Abcam | ab102859 |
| PE / R-Phycoerythrin Conjugation Kit | Abcam | ab102918 |
| Anti-human HLA-A,B,C PE | Biolegend | 311405 |
| Anti-mouse H-2Kb/H-2Db APC | Biolegend | 114613 |
| Anti-mouse H-2Kb/H-2Db PE | Biolegend | 114608 |
| Anti-mouse H-2Kq Alexa Fluor® 647 | Biolegend | 115106 |
| Anti-mouse H-2Kb bound to SIINFEKL APC | Biolegend | 141605 |
| Anti-mCherry Antibody | Rockland | 600-401-P16S |
| Anti-Ki67 (D3B5) Rabbit mAb | Cell Signaling Technology | 9129 |
| DDK (FLAG) mouse monoclonal antibody | Origene | TA180144 |
| Anti-Ki-67 Antibody (Ki-S5) | Sigma-Aldrich | MAB4190 |
| BHLHE41 Polyclonal Antibody | Invitrogen | PA5-25651 |
| Anti-HLA-ABC Monoclonal Antibody (W6/32) | Invitrogen | 14-9983-82 |
| TRA-1-60 antibody | Novus Biologicals | NB100-730 |
| Anti-mouse CD8 | Abcam | ab217344 |
| Anti-mouse CD4 | Abcam | ab183685 |
| Anti-CA19-9 mouse monoclonal antibody | US Biological | Clone 9L426 |
| Sulfo-Cyanine3 NHS ester | Lumiprobe | 41320 |
| Sulfo-Cyanine5 NHS ester | Lumiprobe | 43320 |
| Rhodamine (TRITC) AffiniPure™ Goat Anti-Mouse IgG (H+L) | Jackson Laboratories | 115-025-003 |
| Cy™5 AffiniPure™ Goat Anti-Mouse IgG (H+L) | Jackson Laboratories | 115-175-166 |
| Horse biotinylated anti-rabbit IgG (H+L) | Vector Laboratories | BA-1100 |
| Goat anti-Mouse IgG (H+L) Cross-Adsorbed Secondary Antibody, Alexa Fluor™ 488 | Invitrogen | A-11001 |
| Goat anti-Rabbit IgG (H+L) Cross-Adsorbed Secondary Antibody, Alexa Fluor™ 594 | Invitrogen | A-11002 |
| Goat anti-Rabbit IgG (H+L) Cross-Adsorbed Secondary Antibody, Alexa Fluor™ 488 | Invitrogen | A-11008 |
| Anti-SHARP1 antibody | MyBioSource | MBS8230653 |
| Anti-mouse MHC class I antibody (R1-21.2) | Abcam | ab281904 |
| β2-microglobulin (E8G2O) Rabbit mAb | Cell Signaling Technology | 64046 |
| Anti-luciferase goat polyclonal antibody | Rockland | 200-101-C44S |
| Monoclonal Anti-β-Actin mouse antibody | Sigma-Aldrich | A5441 |
| Anti-β-Tubulin antibody | Abcam | ab6046 |
| Chemicals, peptides, and recombinant proteins |  |  |
| OVA (257-264) Peptide Fragment | Anaspec | AS-60193-1 |
| Experimental models: Cell lines |  |  |
| Ink4a.1 | Gift from Eric Collisson | N/A |
| 6419c5 | Kerafast | EUP005-FP |
| A2441 | Gift from Aram Hezel | N/A |
| A1925 | Gift from Aram Hezel | N/A |
| BxPC-3 | Gift from Arnold Levine | N/A |

| Experimental models: Organisms/strains |  |  |
| --- | --- | --- |
| FVB/NJ | The Jackson Laboratory/Generated | RRID:IMSR_JAX:001800 |
| C57BL/6J | The Jackson Laboratory/Generated | RRID:IMSR_JAX:000664 |
| NU/J | The Jackson Laboratory | RRID:IMSR_JAX:002019 |

## EXPERIMENTAL MODEL AND SUBJECT DETAILS

### Mice

All animal protocols were approved by Rutgers Biomedical and Health Sciences Animal Care and Use Committee and the University of Rochester University Committee on Animal Resources (UCAR). FVB mice, C57Bl/6J and NU/J were purchased from The Jackson Laboratory (Bar Harbor, ME), or bred in-house following established vivarium approved protocols. Young mice (6-8 weeks old) were used for experiment. Mice were monitored daily for survival endpoints. Littermates of the same sex were randomly assigned to experimental groups, except for FVB that only male FVB were used, given that the cell line (i.e. Ink4a.1) implanted is derived from male.

### Patient Samples

Human pancreatic, metastatic, and unaffected specimens from decedents who have previously been diagnosed with pancreatic ductal adenocarcinoma were obtained from the University of Nebraska Medical Center’s Tissue Bank through the Rapid Autopsy Program (RAP) in compliance with IRB 091-01 (Van Andel Institute). Non-cancer tissues are collected in a manner similar to RAP specimens through the UNMC Normal Organ Recovery (NORs) Program in association with LiveOn Nebraska. To ensure specimen quality, organs were harvested within three hours post-mortem and the specimens flash frozen in liquid nitrogen or placed in formalin for immediate fixation. Sections are cut from paraffin blocks of formalin fixed tissue into 4-micron thick sections and mounted on charged slides.

### Cell lines

Ink4a.1 parental (gift from Eric Collisson) is a mouse pancreatic cancer cell line derived from the *p48*^cre^; *Kras*^LSL_G12D^; p16^−/−^/*p19*^−/−^ mouse model (Bardeesy et al., 2006). The cell line was engineered to express luciferase and mcherry (Luc^+^; mCherry^+^) using lentivirus LVP441-PBS (GenTarget). 6419c5 is a KPCY mouse pancreatic cancer cell line purchased from Kerafast. A2441 and A1925 were gifts from Aram Hezel and are murine cell lines derived from pancreatic tumors from Ptf1-cre LSL-Kras^G12D^ p53^fl/+^ Arid1a^fl/fl^ mice. BxPC-3 (gift from Arnold Levine) is a human pancreatic cancer cell line and was authenticated by STR analysis (ATCC). 293T is a human embryonic kidney cell line purchased from ATCC. All cell lines were tested negative for mycoplasma using MycoFluor™ Mycoplasma Detection Kit (Invitrogen). Cells were cultured in complete DMEM media (DMEM (Gibco)+10% FBS+ 1% penicillin/ streptomycin).

## METHOD DETAILS

### Implantation of Tumor Cells

For orthotopic injections, 100 Ink4a.1 Luc^+^mCherry^+^ or 500 6419c5 cells were prepared in a mixture of 10 μL Matrigel Matrix (Corning) and complete DMEM media (1:1). For each mouse, 10 μL of cells were injected into the tail of the pancreas with a microliter syringe and 33-gauge needle (both Hamilton). For experiments in which T cells were quantitated, higher amounts of Dec2KO cell lines (5000 cells of Ink4a *Δ*Dec2 or 6419c5 *Δ*Dec2) had to be injected to generate tumors of an assayable size. The lower gastric vessel was clamped and cut proximal to the spleen. For distal pancreatectomy with splenectomy surgery, an incision was made in the abdomen to expose the primary pancreatic tumor and spleen. The lower gastric vessel was clamped and cut proximal to the spleen. The healthy pancreas was clipped and cut. The primary tumor and spleen were removed *en bloc* 3-4 weeks after implantation. Intrasplenic injections were performed as previously described(Dudgeon et al., 2023), and liver was harvested 3 weeks after intrasplenic injection. Buprenorphine and bupivacaine were given for pain management. Immediately following surgery, mice were allowed to recover under a heating lamp until ambulatory.

### MRI and IVIS Imaging

Mice with a DFI>300 days were sent to the Rutgers Molecular Imaging Center and MR images acquired with a 1T M2-High Performance MRI System (Aspect Magnet Technologies). Mice that had been orthotopically injected with Ink4a.1 cells were injected with 15 mg/kg luciferin (Pierce) and imaged using the IVIS Spectrum (Caliper Life Sciences). LivingImage software (Revvity) was used to set the minimum and maximum luminescent rate to be equal in all images.

### Isolation of Tumor cells and Liver Disseminated Tumor Cells (DTCs)

Primary tumor, early liver metastasis and liver tissues from dormant mice (disease free interval >120 days) were minced with razor blades and enzymatically digested in DMEM+10% FBS with 200 units/mL collagenase type I, 60 units/mL hyaluronidase, and 50 μg/mL DNase I at 37°C for 45 minutes, with vortexing every 5 minutes. Tissue pieces were mashed through a 100 μm cell strainer (PluriSelect) and spun at 500 xg for 5 minutes. For liver samples, red blood cells were lysed using ACK Lysing Buffer (Gibco) for 3 minutes, and then washed once with cold PBS. Cells were counted using TC20 Automated Cell Counter (BioRad). 1 million cells/sample were resuspended in FACS buffer (1x DPBS/2% BSA) containing 1x DAPI and sorted for mCherry^+^, DAPI^-^ expression on BD Biosciences Influx High Speed Cell Sorter.

### Flow Cytometry Analysis of Dormant DTCs

Liver cell suspensions were prepared as described above, while cultured cell lines were prepared by gentle trypsinolysis. One million cells were resuspended in 100 μl FACS buffer and blocked with TruStain FcX antibody (Biolegend) at 100x dilution at 4 °C for 10 minutes. After washing and spinning, each sample was resuspended with 100 μl antibody cocktail containing anti-mouse CD44 AF700 (Biolegend), anti-mouse CD133 PE-Cy7 (Biolegend) antibodies and incubated at 4 °C for 30 minutes in the dark. Samples were washed with FACS buffer, fixed and permeabilized using the True-Nuclear Transcription Factor Buffer Set (Biolegend) according to manufacturer’s instructions. For cyclin-dependent kinases staining, samples were further stained with anti-p21 AF488 and anti-p27 PE (both Cell Signaling Technology). For Dec2 staining, anti-SHARP1 primary antibody (Abcam) was conjugated to PE using the PE/R-Phycoerythrin Conjugation Kit - Lightning-Link (Abcam) according to the manufacturer’s instructions. Each sample was stained with 5 μL of conjugated anti-SHARP1 PE and anti-Ki67 APC-Cy7(Biolgend) at 4 °C for 45 minutes in the dark, washed with 1x permeabilization buffer and resuspended in 300 μL FACS buffer prior to flow cytometry. DTCs were classified as CD44^+^/CD133^+^/mCherry^+^. The ABC total compensation bead kit (Invitrogen) was used according to the manufacturer’s instructions to determine compensation for multicolor flow cytometry.

### Nucleic Acid Extraction

Genomic DNA was extracted using DNAzol (Invitrogen) for sorted DTCs and Wizard Genomic DNA Purification kit (Promega) for standard samples according to manufacturer’s instructions. Plasmid DNA was extracted using QIAprep Spin Miniprep Kit (Qiagen) according to manufacturer’s instructions. RNA was extracted using RNeasy kit (Qiagen) was used for standard RNA and RNAseq samples, while Quick RNA miniprep kit (Zymo Research) was used for mouse tails according to manufacturer’s instructions.

### Ultra-Low RNA-seq Analysis

DTCs were profiled using ultralow input RNA-seq protocol while primary tumor and metastatic cells were profiled using standard RNA-seq. Transcript expression was calculated using the log2 values of the significant non-zero values of TPMs. Primary tumor and metastatic cells were profiled using standard RNAseq. After quality check using FastQC (Babraham Bioinformatics)(Brown et al., 2017), Salmon (v.1.1.0) was used to quantify transcript-level expression and EdgeR (v.3.31.4) to identify genes with significantly differential expression between pairs of conditions based on replicated count data from bulk RNA-seq profiling(Liao et al., 2019). The p-values were corrected for multiple testing using FDR. Heat maps were created using Morpheus (Broad Institute, https://software.broadinstitute.org/morpheus).

### Transfection and Immunocytochemistry (ICC)

Ink4a.1 cells were plated onto glass coverslips and the next day transfected with the expression plasmid pCMV6-mouse Bhlhe41-DDK-myc (Origene). Two days later, cells were fixed with 4% paraformaldehyde at room temperature for 10 minutes and permeabilized with 0.25% Triton for 10 minutes. Cells were incubated 4 °C overnight with anti-DDK (Origene) and anti-mouse Ki67 (Cell Signaling Technology) primary antibodies. Anti-mouse AF488 and anti-rabbit AF594 secondary antibodies (both Invitrogen) were used to detect DDK (i.e. Dec2) and Ki67, respectively. For MHC-I staining, cells were incubated with anti-mouse MHC-I primary antibody (Abcam) at 4 °C overnight and with anti-rabbit AF488 the next day. Coverslips were mounted using Vectashield plus DAPI (Vectorlabs) and images taken using a Keyence BZ-X710 fluorescent microscope.

### PCR/quantitative-PCR (qPCR) on DTCs

Genomic DNA (gDNA) was extracted as described above. Luciferase presence/absence in gDNA was measured using TaqMan Universal PCR Master Mix No AmpErase UNG (Applied Biosystems) and commercial Taqman Gene Expression Assays for luciferase (ThermoFisher Scientific) on a 7500 Real Time PCR machine (Applied Biosystems). For *Kras* PCR, RNA was extracted as mentioned above and reverse transcribed into cDNA using the Reverse Transcription Kit (Applied Biosystems) according to manufacturer’s instructions. Primers for WT (Forward: 5’-ACT-TGT-GGT-GGT-TGG-AGC-TGG-3’; Reverse: 5’-CGT-AGG-GTC-ATA-CTC-ATC-CAC-A-3’) and *Kras*^G12D^ mutant (Forward: 5’-ACT-TGT-GGT-GGT-TGG-AGC-AGA-3’; Reverse: same as WT reverse) were used for PCR using Platinum™ Taq DNA Polymerase (Invitrogen), with 35 cycles of 95°C for 1 minute, 58°C for 30 seconds and 72°C for 20 seconds on SimpliAmp Thermo Cycler (Applied Biosystems). PCR products were visualized on a 2% TAE agarose gel stained with ethidium bromide.

### Multiplexed Immunofluorescence (IF)

We performed immunofluorescence on 5 µm thick sections cut from formalin-fixed, paraffin-embedded (FFPE) blocks. Paraffin was removed from 5 µm thick FFPE sections using CitriSolv Hybrid (Decon Labs) containing d-limonene and isopropyl cumene, and the tissue was rehydrated through an ethanol gradient of 100%, 95%, and 70% followed by washing with PBS. Following rehydration, antigen retrieval was achieved through incubating slides in citrate buffer at 100 °C for 20 minutes. Slides were blocked in phosphate-buffered saline with 0.05% Tween-20 (PBST 0.05) and 3% bovine serum albumin (BSA) for 1 hour at room temperature (RT). Primary anti-KI-67 antibody (Sigma-Aldrich), anti-BHLHE41 polyclonal antibody and anti-HLA class1 polyclonal antibody (both Invitrogen) were diluted in blocking buffer (1:250) and applied to sections for 2 hours or at 4 °C overnight. Cy5-conjugated anti-mouse, or TRITC-conjugated anti-rabbit secondary antibodies (The Jackson Laboratory) were diluted in blocking buffer (1:500) and incubated with sections for 2 hours. Primary antibodies against CA19-9 (clone 9L426, US Biological Life Sciences), and TRA-1-60 (Novus Biologicals) were labeled for immunofluorescent staining with Sulfo-Cyanine5 NHS ester and Sulfo-Cyanine3 NHS ester respectively (both Lumiprobe). After dialysis to remove unreacted conjugate, the antibodies were diluted into the same solution of PBST 0.05 with 3% BSA to a final concentration of 10 µg/mL. Slides were incubated overnight with this solution at 4 °C in a humidified chamber.

The following day, the solutions were decanted, and the slides were washed twice in PBST 0.05 and once in 1X PBS, each time for 3 minutes. The slides were dried via blotting and incubated with DAPI at 10 µg/mL in 1X PBS for 15 minutes at RT. Two five-minute washes were performed in 1X PBS, and then slides were cover-slipped and scanned using AxioScan.Z1 microscope (Zeiss). We next quenched the fluorescence using 6% H_2_O_2_ in 250 mM sodium bicarbonate (pH 9.5-10) and performed another round of immuno-fluorescence using two different antibodies. The subsequent incubations and scanning steps were as described above.

Prior to the second round of detection with the TRA-1-60 antibody, we treated the slides with sialidase to remove terminal sialic acids. The slides were incubated with a 1:200 dilution (from a 50,000 U/mL stock) of α2-3,6,8 Neuraminidase in 5 mM CaCl_2_, 50mM pH 5.5 sodium acetate overnight at 37 °C. The subsequent incubations and scanning steps were as described above. In each round of fluorescence imaging, the microscope collected 3 images at each field-of-view, each image corresponding to the emission maxima of DAPI, Cy3, and Cy5. The three images were saved as independent, stacked layers in a single file. Following all fluorescence imaging, the slides were stained with hematoxylin and eosin (H&E) using a standard protocol and imaged by brightfield microscopy using the Aperio ScanScope (Leica Biosystems).

### Image Analysis

All fluorescence images were initially processed using SignalFinder(Barnett et al., 2019). SignalFinder automatically creates a map of the locations of pixels containing signal in each layer of the multiplexed immunofluorescence. This map was aligned to the H&E image based on the signals from cell nuclei using the Warpy extension in QuPath(Bankhead et al., 2017). We automated cell nuclei detection followed by a 10 µm expansion to estimate cell borders and then quantified the percentage of positive glycan pixels in each cell for each image layer.

We developed the SignalFinder software using MATLAB, Java, and C++. We used Microsoft Excel and MATLAB for analyzing numerical output; GraphPad Prism, the R language, and Microsoft PowerPoint for the preparation of graphs, and Canvas XIV for the preparation of figures.

### Immune Depletion

CD4 or CD8+ T cells were depleted using *α*-CD4 and *α*-CD8 InVivo MAb (both Bio X Cell). Mice were I.P. injected with 100 μL of 2.5mg/ml antibody for 3 days prior to tumor cell implantation, then once every 4 days after tumor implantation for the duration of the experiment.

### Bulk RNA-seq Analysis

RNA was extracted as described above for bulk RNAseq. For demultiplexing, raw reads generated from the Illumina base calls were demultiplexed using bcl2fastq (v.2.20.0). Quality filtering and adapter removal are performed using FastP (v.0.23.1) with the following parameters: "--length_required 35 -- cut_front_window_size 1 --cut_front_mean_quality 13 --cut_front --cut_tail_window_size 1 -- cut_tail_mean_quality 13 --cut_tail -y -r" ^14^. For quality control, processed reads were then mapped to GRCm39/gencode M31 using STAR_2.7.9a with the following parameters: "--twopass Mode Basic -- runMode alignReads --outSAMtype BAM Unsorted -- outSAMstrandField intronMotif -- outFilterIntronMotifs RemoveNoncanonical --outReadsUnmapped Fastx" ^15^. For alignment, gene level read quantification was derived using the subread-2.0.1 package (featureCounts) with a GTF annotation file GRCm39/gencode M31 and the following parameters for stranded RNA libraries "-s 2 - t exon -g gene_name"(Chen et al., 2018; Frankish et al., 2021). Differential expression analysis was performed using DESeq2-1.34.0 with a P-value threshold of 0.05 within R (v.4.0.2, https://www.R-project.org/). Principle component analysis plot was created within R using pcaExplorer c2.14.2 to measure sample expression variance. Heatmaps were generated using the pheatmap (v.1.0.12) and were given the rLog transformed expression values. Gene ontology analyses were performed using the EnrichR package. Volcano plots and dot plots were created using ggplot2.

### Quantitative PCR (qPCR) Analysis of Cell Lines

cDNA was made using the TaqMan Reverse Transcription Reactions (Applied Biosystems) according to manufacturer’s instructions. The presence of luciferase gene was measured using a customized TaqMan gene expression assay (5’-GCT-GAT-GCC-CAT-GCT-GTT-C-3’; 5’-TGT-TGG-GTG-CCC-TGT-TCA-TC-3’; 5’-FAM-CCA-GCT-AAC-GAC-ATC-TAC-A-NFQ-3’). All other genes were measured using commercial Taqman Gene Expression Assays (ThermoFisher Scientific), except for TBP which was measured using SYBR reagents and oligonucleotides 5’-CCA-GAA-CTG-AAA-ATC-AAC-GCA-G-3’ and 5’TCT-ATC-TAC-CGT-GAA-TCT-TGG-C-3’. qPCR assays were performed on a 7500 Real Time PCR System using either TaqMan Universal PCR Master Mix No AmpErase UNG, or PowerUp SYBR Green Master Mix (both ThermoFisher Scientific). Gene expression level was normalized with human or mouse β-actin (Applied Biosystems) except in circadian experiments, where b-actin was inappropriate and TBP, PPIA or RPLPO were used as substitutes as described in figures.

### CUT&RUN Sequencing Analysis

To assess the genomic distribution of modified histones, CUT&Tag was performed using the Epicypher® CUTANA™ Direct-to-PCR CUT&Tag Protocol (v.1.7). Briefly, nuclei isolated from 100,000 cells were incubated with Concanavalin-A beads (Bangs Laboratories BP531) followed by the addition of primary antibody (H3K27Ac, Abcam 4729; H3K27me3, Cell Signaling Technology 9733) and incubation overnight at 4°C. Nuclei bound beads were resuspended in buffer containing antirabbit secondary antibody (Antibodies Online Inc. ABIN101961) and incubated for an additional for 30min at RT. CUTANA™ pAG-Tn5 preloaded with Illumina sequencing adapters was added for 10 min at room temperature, followed by activation with MgCl_2_ and tagmentation at 37°C for 1hr. The reaction was stopped and DNA was isolated by successive incubation with TAPS Buffer, SDS Release Buffer, SDS Quench Buffer. The library was amplified with uniquely barcoded primers with NEBNext High Fidelity 2x PCR Master Mix (NEB M0541) and purified with 1.3x volume Agencourt AMPure XP Magnetic Beads (Beckman Coulter A63880). Assays were performed in duplicate for each antibody and each cell line. For Dec2 genomic distribution, CUT&RUN was performed using the EpiCypher® CUTANA™ CUT&RUN Protocol v2.0. Cells (500,000) were permeabilized with digitonin, bound to Concanavalin-A beads (Bangs Laboratories BP531) and then incubated with primary antibody (Sharp 1, MyBioSource, MBS8230653) overnight at 4°C. CUTANA™ pAG-MNase was added at room temperature for 10 min followed by the addition of 100mM CaCl_2_ to activate cleavage for 2hr at 4°C. MNase activity was halted with Stop Buffer for 10min at 37°C. DNA was purified using the Qiagen MiniElute Kit (Qiagen 28004). Reactions were performed in duplicate and library preparation was performed using Illumina adapters and NEBNext Ultra II DNA Kit.

Sequencing was performed on the Illumina NovaSeq 6000 platform (50bp PE) by the University of Rochester Genomics Research Center. Raw sequencing data was subjected to quality trimming using fastp (v.b1) and aligned to the GRCm39 reference genome (NCBI assembly GCF_000001635.27) using the bowtie2 package (v.2.3.5.1) and a mapping quality threshold MAPQ< 10. Duplicate aligned reads were removed using Picard (v.2.12.0, http://broadinstitute.github.io/picard). BAM files were converted to BigWig files and RPKM normalized using DeepTools (v.2.5.3) and visualized in IGV(Ramirez et al., 2014).

### Proteasome Activity

Proteasome activity was assayed in freeze-thaw-generated lysates prepared from cultured cells as previously described(Javitt et al., 2023), with the following modifications: To normalize, protein concentrations were assayed using Pierce BCA kit (ThermoFisher Scientific) using a Synergy 2 spectrophotometer (BioTek). Peptide substrate Suc-Leu-Leu-Val-Tyr-AMC (Suc-LLVY-AMC) (Bachem) was added at 10 µM to 10 µg of total protein. Fluorescence increase resulting from degradation of peptide-AMC at 37 °C was monitored over time by means of a fluorometer (Synergy 2 Hybrid Multi-Mode Microplate Reader, BioTek) at 360-nm excitation and at 460-nm emission. Pancreatic cell lines have residual chymotrypsin activity, which had to be subtracted in order to generate the actual proteasome activity. Residual chymotrypsin activity was determined by re-assaying in the presence of 100 nM of the proteasome-specific inhibitor MG132.

### Generation of Overexpressing, Knockout or Knockdown Cell Lines

CRISPR-Cas9-directed knockout of Dec2 in cell lines 6419c5 and Ink4a was performed using a commercial lentivirus particle purchased from Horizon Discovery (VSGM11942-248549844 or VSGM11942-248123446). Lentivirus particles for CRISPR-dCas9-directed knockdown of b2m were generated after subcloning a 20 nt sequence (5’ GCC-GTC-GGG-AAG-GAG-AAC-TA-3’ against the 5’- UTR of b2m into pLV-hU6-sgRNA hUBC-dCas9-KRAB-T2a-Puro (gift from Charles Gersbach, Addgene plasmid # 71236; http://n2t.net/addgene:71236; RRID: Addgene_71236). For some constructs, pLV-hU6-sgRNA hUBC-dCas9-KRAG-T2a-bsd with stuffer (a gift from Stephano Mello) was used instead of pLV-hU6-sgRNA hUBC-dCas9-KRAB-T2a-puro. Overexpression of b2m was performed by transfecting cells with lentivirus particles generated after subcloning the open reading frame for b2m into plasmid pCDH-EF1-FHC, which was a gift from Richard Wood (Addgene plasmid # 64874; http://n2t.net/addgene:64874; RRID: Addgene_64874). Lentiviral particles were generated by co-transfecting 293T cells with plasmid of interest along with pMD2.g and psPAX2, as recommended by Addgene. pMD2.G and psPAX2 (gifts from Didier Trono, Addgene plasmid # 12259; http://n2t.net/addgene:12259; RRID: Addgene_12259 and Addgene plasmid #12260; http://n2t.net/addgene:12260; RRID: Addgene_12260). Transfection of pancreatic cell lines and selection of overexpressers, knockdowns and knockouts were performed as recommended by Addgene. Prior to transfection, individual cells within cell line cultures were “cloned” by limiting dilution, and these clones were used for transfection. Cells were then transfected with lentivirus particles that either expressed the gene or sgRNA of interest, or else empty vector. For knockouts, individual clones were again collected by limiting dilution, then screened for loss of expression of Dec2 using qPCR and Western blot. Both alleles were confirmed to be knocked out by DNA sequencing of 20 TOPO-TA clones generated by subcloning genomic PCRs that flanked the sgRNA target sequences. Knockdowns and overexpressers were not individually cloned and instead represent pools of transfected cells.

### Flow Cytometry Analysis of Cell Lines

Adherent pancreatic cell lines were removed from plates by gentle trypsinization for 120 seconds, then washed extensively with 10% serum to inactivate trypsin and then twice with FACS buffer. Cells were then blocked with anti-mouse CD16/CD32 Fc block (1:100, BD Biosciences) in FACS buffer for 10 minutes at 4°C and stained with antibodies diluted in FACS buffer for 30 minutes at 4°C in the dark. After staining, cells were washed and resuspended in FACS buffer. To detect intracellular proteins, cells were fixed and permeabilized using True Nuclear Transcription Factor Buffer Set (Biolegend) according to manufacturer’s instructions, stained with antibodies diluted in 1x permeabilization buffer from the kit for 45 minutes at 4°C in the dark. Finally, cells were washed with 1x permeabilization buffer and resuspended in FACS buffer and analyzed on BD LSR Fortessa Cell Analyzer (BD Biosciences).

For circadian experiments, 300 ng/ul of the peptide SIINFEKL (AnaSpec) was added to one million gently trypsinized, circadian-synchronized cells in FACS buffer for 10 mins on ice, then washed twice in FACS buffer. Cells were then treated with antibody that detects SIINFEKL bound to H-2Kb (Biolegend). After 30 minutes with antibody at room temperature in the dark, cells were washed twice with FACS Buffer, then put on ice prior to flow cytometry analysis.

### Immunohistochemistry (IHC) Staining and Quantification

Primary tumors or livers were fixed in 10% formalin (VWR) and embedded in paraffin before trimmed and cut into slides. Slides were deparaffined in Histoclear (National Diagnostics) for 5 minutes for a total of 3 times. Slides were rehydrated through 2×100% ethanol, 2×95% ethanol, 1×70% ethanol, 1×50% ethanol and 2xdH_2_O. Antigens were retrieved by boiling the slides for 15 minutes in Citrate-Based Antigen Unmasking Solution (Vector Laboratories). Slides were blocked in BLOXALL Endogenous Blocking Solution (Vector Laboratories) for 10 minutes and washed 3x TBST buffer (1xPBS 1% Tween). Endogenous enzyme activity was blocked by Streptavidin/Biotin Blocking Kit (Vector Laboratories) for 30 minutes each. Slides were blocked using 10% normal serum of appropriate species for 30 minutes and then stained with primary antibodies overnight at 4 °C. Slides were washed 3x TBST buffer and stained with biotinylated HRP secondary antibody (Vector Laboratories) for 1 hour at room temperature. Slides were washed 3x TBST buffer, incubated with Vectastain ABC HRP Kits and visualized with ImmPACT DAB Substrate Peroxidase Kit (both Vector Laboratories). Slides were further counterstained with hematoxylin and mounted with Permount Mounting Medium (Fisher Scientific). Slides were scanned using Olympus VS120 Virtual Slide Microscope (Olympus) and positive staining was quantified with Visiopharm Image Analysis System (Visiopharm).

### Cytokines/Chemokines Secretion Profiling

Tumor cell lines were seeded in 6-well plates (Falcon) in 1 ml serum free DMEM media overnight. On the next day, the media was collected and spun at 2,000 xg for 5 minutes at 4 °C. The supernatant was analyzed using Mouse Cytokine Microarray (R&D Systems). The resulting blot was imaged on ChemiDoc MP Imaging System and analyzed with Image Lab software (both Bio-Rad). Quantification of cytokine was determined by pixel density on the blot, normalizing to the density of WT for each detected cytokine.

### Western Blot Analysis

Cells were harvested and lysed using buffer consisting of 1xRIPA, Phosphatase Inhibitor Cocktail and PMSF (both Cell Signaling Technology). Protein concentration was determined using Pierce BCA Assay (Invitrogen) according to manufacturer’s instruction. Equal amount of proteins from each sample were mixed with b-ME/4xLameli buffer (1:100), loaded on Mini-Protean TGX Gels and ran in 1xTGS running buffer (all Bio-Rad). Gel was equilibrated in 1x TBST buffer and transferred to a PVDF membrane in semi-dry transfer buffer. The membrane was block in 0.5% non-fat milk TBST buffer for 1 hour and incubated with primary antibody overnight at 4 °C. On the next day, the membrane was washed 3 times with 1x TBST buffer and incubated with appropriate secondary antibody diluted in blocking buffer for 1 hour. The membrane was washed 3 times with 1x TBST buffer and incubated in Western Lightning Plus-ECL (Revvity Health Sciences) or SuperSignal West Femto Maximum Sensity Substrate (ThermoFisher Scientific Scientific). The membrane was then imaged on ChemiDoc MP Imaging System and analyzed with Image Lab.

### Cell Line Doubling Time Measurements

Tumor cell lines were seeded in 6 well-plates at 2000 cells/well on day 0. Cell confluency was determined twice per day for 5 days using Celigo S Image Cytometer (Revvity) under bright field cell detection.

### Promoter assays

Promoter sequences for the mouse Cdk4 and b2m genes were inserted into the firefly luciferase promoter fusion plasmid pGL4.10 (Promega). The Cdk4 and b2m promoters were obtained from mouse gDNA using oligonucleotides 5’-TGA-GCT-CGC-TAG-CGA-GGC-TAG-CTA-CGT-GTC-CAT-3’ and 5’-AGA-TCT-TGA-TAT-CCC-AGA-CCA-TAG-ACA-CAG-GCC-3’ (Cdk4) or 5’-TGA-GCT-CGC-TAG-CAC-ACA-CAGGG-CTT-TTG-TCT-T-3’ and 5’-AGA-TCT-TGA-TAT-CCT-GAA-CTC-ACA-GCC-AAT-CCG-3’ (b2m). The two PCRs were separately recombined into XhoI-digested pGL4.10 using an InFusion HD kit (Takara). 1.5 μg of each promoter/fusion construct was then transiently transfected for 48 hours into 100,000 Ink4a PDAC cells, along with 1.5 ug of Dec2 expression plasmid (Origene #MR223412) and 10 ng of renilla luciferase-expressing plasmid pRL-SV40 (Promega), using lipofectamine 2000 (ThermoFisher Scientific Scientific). After two days, cells were washed twice with PBS, lysed with passive lysis buffer, then assayed for firefly and renilla luciferases using a dual luciferase kit (Promega).

### Circadian analysis

For SIINFEKL/MHC-1 assays, 400,000 cells were plated onto 10 cm dishes, treated the next day with 10nM forskolin for one hour, then washed twice with PBS, then re-fed with DMEM-10% FBS. 24-64 hours later, cells were gently trypsinized for exactly 120 seconds, then split into different tubes for separate assays of surface MHC-I, SIINFEKL-binding to MHC-I, RNA, and proteasome activity. For RNA analyses, circadian rhythms were established using 10 nM of dexamethasone. 50,000 cells plated onto 6 well dishes were treated the next day with 10 nM dexamethasone, then harvested for RNA 24-72 hours later with 4 hours interval.

### T cell killing assay

A1925 cells were synchronized with 10 nM forskolin for one hour, then washed twice and treated with normal media for 32 or 40 hours. Cells were then treated with SIINFEKL peptide (ova) and seeded in 24-well plates at 20,000 cells/well in triplicates. Effector cells (OT-1 T cells) were added to the target cells at effector to target ratios of 10:1. No ova treated cells were used as negative control. After 24 h of co-culture at 37 °C, 7-AAD dye was added for 10 min, and cells were analyzed on a BD LSR II flow cytometer (BD Biosciences). Unpaired t test was used to determine statistical significance of 7AAD+ tumor cell% between A1925 synchronized for 32 hours and 40 hours.

## QUANTIFICATION AND STATISTICAL ANALYSIS

### Statistical Analysis

Statistical analysis of comparisons between two groups were performed using Student’s unpaired t test or multiple unpaired t test using FDR approach. Significance of positive markers in multiplexed IF staining were identified by mixed effects logistic regression with post hoc comparisons. Significance of overall survival between two groups was determined by Kaplan-Meier analysis using log-rank test. Significance of cell line doubling time was determined by non-linear regression with exponential growth model. All statistical analyses (except for circadian analysis) and quantitative graphs were performed/generated with GraphPad Prism 10 (GraphPad).

### Circadian Analysis

The Extended Circadian Harmonic Oscillator (ECHO) application was used for oscillation analysis of qPCR and proteasome activity data. This application uses parametric approaches to identify rhythmicity in oscillators from biological data. The analysis was performed in ECHO v4.0. The following parameters were used: paired replicates, linear trends removed, and default parameters to determine if oscillation is harmonic, overexpressed, or repressed. mRNA expression or proteasome activity were determined to have circadian rhythmicity if they had a period between 20-26 hours, and had a p < 0.05, and a BH.Adj.P < 0.05.

---

**Table S1.** List of Human PDAC liver metastatic tissues annalzyed by ROI size and quantitated (percent) for glycan staining.

| Tissue | Cluster size |  |  | <100 |  |  | 100-1000 |  |  | >1000 |  |  |
| --- | --- | --- | --- | --- | --- | --- | --- | --- | --- | --- | --- | --- |
|  |  |  |  | CA199 Only Cells | sTRA Only Cells | Dual Expressing Cells | CA199 Only Cells | sTRA Only Cells | Dual Expressing Cells | CA199 Only Cells | sTRA Only Cells | Dual Expressing Cells |
|  | <100 | 100-1000 | >1000 | Average | Average | Average | Average | Average | Average | Average | Average | Average |
| 118-RF-4736 | 13 | 11 | 6 | 49.20% | 0.90% | 0.00% | 27.60% | 0.40% | 0.60% | 31.70% | 1.10% | 0.90% |
| 123-RF-4934 | 29 | 18 | 5 | 14.40% | 45.80% | 6.10% | 8.20% | 36.70% | 4.10% | 0.30% | 68.80% | 1.10% |
| 156-RF-6503 | 1 | 3 | 1 | 80.90% | 0.00% | 0.00% | 53.00% | 0.10% | 1.60% | 53.10% | 0.10% | 0.70% |
| 44-RF-1796 | 2 | 15 | 2 | 24.40% | 18.60% | 2.30% | 19.70% | 21.20% | 2.30% | 0.80% | 57.50% | 1.00% |
| 45-RF-1826 | 1 | 3 | 2 | 0.00% | 16.20% | 0.00% | 8.30% | 15.10% | 15.40% | 33.00% | 3.30% | 34.10% |
| 54-RF-2156 | 2 | 5 | 5 | 39.20% | 5.00% | 30.80% | 16.00% | 6.10% | 10.40% | 9.70% | 6.70% | 10.00% |
| 58-RF-2307 | 2 | 6 | 3 | 71.90% | 0.00% | 0.00% | 23.00% | 11.50% | 14.20% | 41.80% | 1.30% | 11.90% |
| 66-RF-2656 | 8 | 9 | 1 | 0.00% | 58.60% | 0.00% | 4.10% | 52.00% | 3.40% | 0.00% | 67.10% | 0.10% |
| 77-RF-3143B | 14 | 18 | 7 | 11.70% | 31.70% | 10.90% | 28.00% | 20.50% | 10.00% | 17.50% | 16.40% | 9.70% |
| 85-RF-3448 | 20 | 41 | 8 | 31.70% | 0.20% | 0.20% | 27.40% | 0.10% | 0.40% | 32.80% | 1.40% | 1.30% |

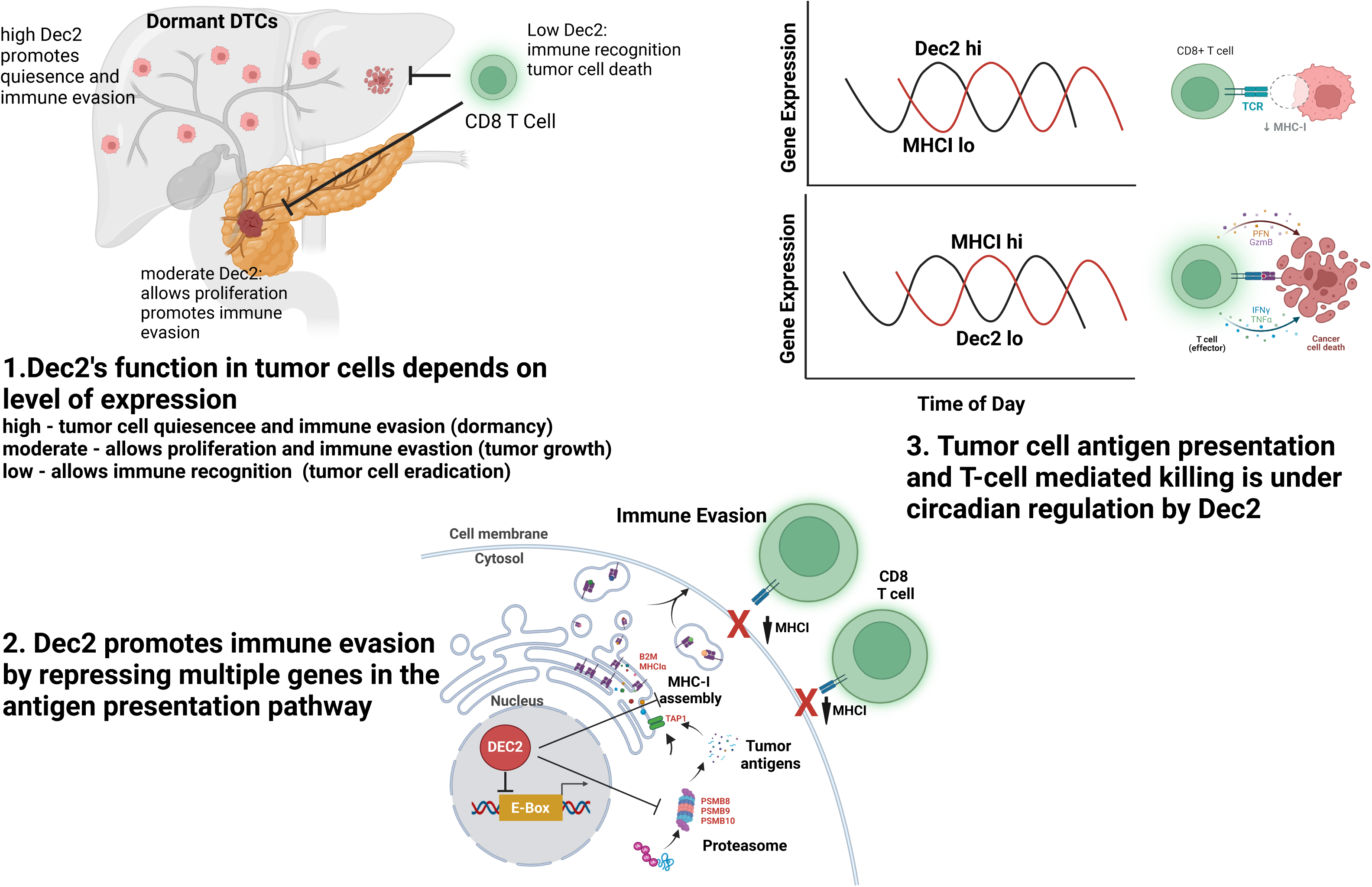

