## Supplementary figures and images for "The circadian gene Dec2 promotes pancreatic cancer dormancy by regulating tumor cell antigen presentation to facilitate immune evasion"

### Image_001.png

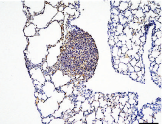

### Image_002.jpg

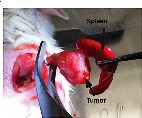

### Image_003.jpg

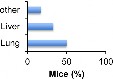

### Image_004.jpg

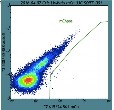

### Image_005.jpg

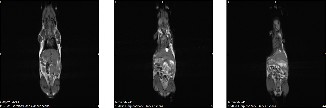

### Image_006.jpg

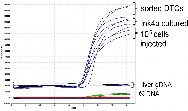

### Image_007.jpg

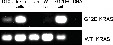

### Image_008.png

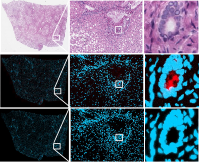

### Image_009.png

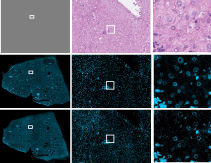

### Image_010.png

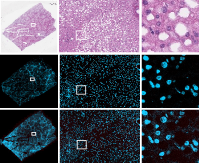

### Image_011.png

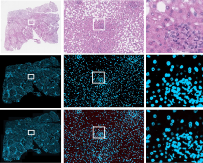

### Image_012.png

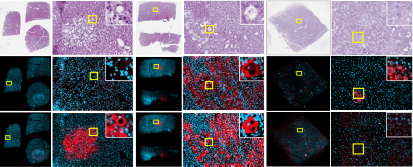

### Image_013.png

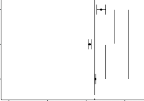

### Image_014.png

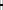

### Image_015.png

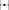

### Image_016.png

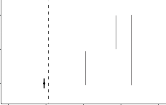

### Image_017.jpg

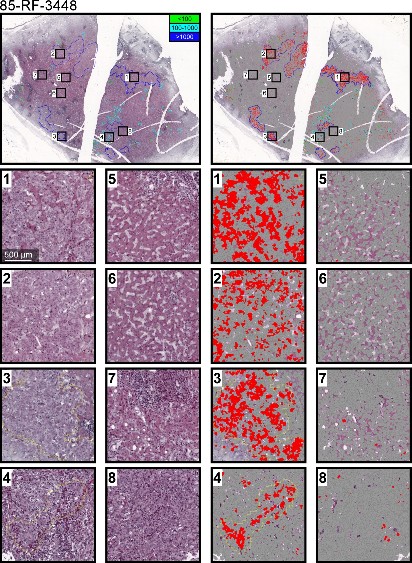

### Image_018.png

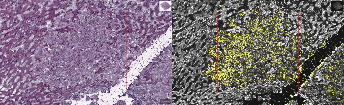

### Image_019.png

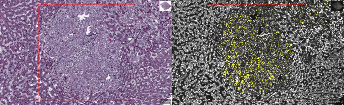

### Image_020.png

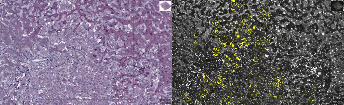

### Image_021.png

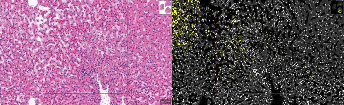

### Image_022.png

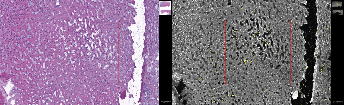

### Image_023.png

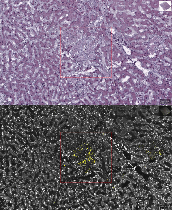

### Image_024.png

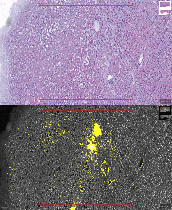

### Image_025.png

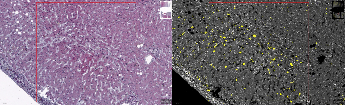

### Image_026.png

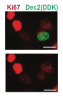

### Image_027.png

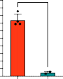

### Image_028.png

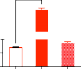

### Image_029.png

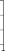

### Image_030.png

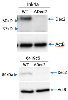

### Image_031.png

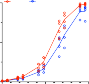

### Image_032.png

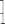

### Image_033.png

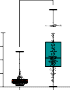

### Image_034.png

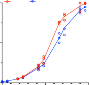

### Image_035.png

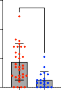

### Image_036.png

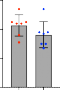

### Image_037.png

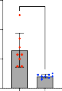

### Image_038.png

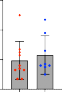

### Image_039.png

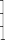

### Image_040.png

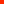

### Image_041.png

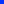

### Image_042.png

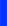

### Image_043.png

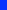

### Image_044.png

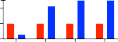

### Image_045.png

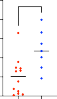

### Image_046.png

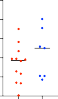

### Image_047.png

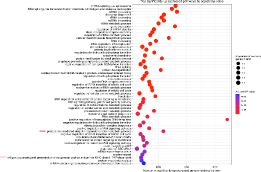

### Image_049.png

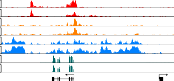

### Image_050.png

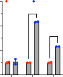

### Image_051.png

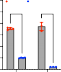

### Image_052.png

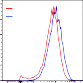

### Image_053.jpg

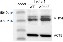

### Image_055.jpg

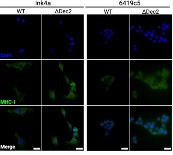

### Image_056.png

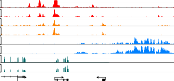

### Image_058.png

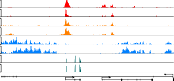

### Image_059.jpg

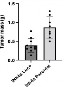

### Image_060.png

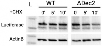

### Image_062.png

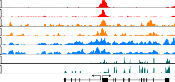

### Image_063.png

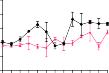

### Image_064.png

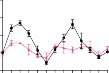

### Image_065.png

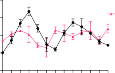

### Image_066.png

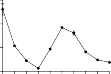

### Image_067.png

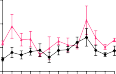

### Image_068.png

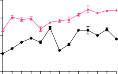

### Image_069.png

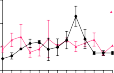

### Image_070.png

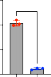
